# Natural killer (NK) cell-derived extracellular-vesicle shuttled microRNAs control T cell responses

**DOI:** 10.1101/2022.01.05.475119

**Authors:** Sara G. Dosil, Sheila López-Cobo, Ana Rodríguez-Galán, Irene Fernández-Delgado, Marta Ramirez-Huesca, Paula Milán-Rois, Milagros Castellanos, Álvaro Somoza, Manuel José Gómez, Hugh T. Reyburn, Mar Valés-Gómez, Francisco Sánchez-Madrid, Lola Fernández-Messina

## Abstract

Natural killer (NK) cells recognise and kill target cells undergoing different types of stress. NK cells are also capable of modulating immune responses. In particular, they regulate T cell functions. Small RNA next-generation sequencing of resting and activated NK cells and their secreted EVs led to the identification of a specific repertoire of NK-EV-associated microRNAs and their post-transcriptional modifications signature. Several microRNAs of NK-EVs, namely miR-10b-5p, miR-92a-3p and miR-155-5p, specifically target molecules involved in Th1 responses. NK-EVs promote the downregulation of GATA-3 mRNA in CD4^+^ T cells and subsequent T-bet de-repression that leads to Th1 polarization and IFN-γ and IL-2 production. NK-EVs also have an effect on monocyte and moDCs function, driving their activation and increased presentation and co-stimulatory functions. Nanoparticle-delivered NK-EV microRNAs partially recapitulate NK-EV effects *in vivo*. Our results provide new insights on the immunomodulatory roles of NK-EVs that may help to improve their use as immunotherapeutic tools.

## INTRODUCTION

Extracellular vesicles (EVs) are key mediators of cell-to-cell communication and play a crucial role in the regulation of immune responses (1). EVs, e.g. exosomes, microvesicles and apoptotic bodies, carry bioactive molecules such as proteins, carbohydrates, lipids, but also genetic information, including microRNAs (miRNAs) (2). These miRNAs can be transferred among cells and modulate gene expression in the recipient cell.

Natural killer (NK) cells recognise and kill cells undergoing different types of stress, including aging, malignant transformation and pathogenic infection. NK cells also modulate immune responses, particularly T cell function. NK cells promote T cell differentiation, proliferation and cytokine production (3). These effects are mediated by direct interactions between NK and T lymphocytes; but also indirectly through their effect on antigen-presenting cells and the secretion of soluble factors.

Every immune cell releases EVs, including NK cells. Recent evidence shows that NK-derived EVs (NK-EVs) can exert antitumoral functions. NK cell-derived exosomes mediate cytolytic effects on melanoma (4) and neuroblastoma cells (5). In addition, NK cell-derived exosomes contain cytotoxic molecules (6), such as Fas-L and perforin, which induce tumour cell apoptosis (7); or DNAM1 (8), which is also involved in exosome-mediated antitumoral responses, as revealed by experiments using blocking antibodies. Proteomic analyses of NK-EVs have identified additional effector candidates that may induce tumour cell death, including TRAIL, NKG2D or fibrinogen (9). Thus, increasing evidence supports the potential use of cytotoxic cell-derived EVs, including NK cells and also cytotoxic T lymphocytes (CTLs) as therapeutic agents (10). Although most studies to date have highlighted the potential role of NK-EVs in cancer, they can be potentially used to modulate other biological processes and pathologies. In particular, NK-EVs have a beneficial effect in lung injury recovery after *Pseudomonas aeruginosa* infection (11). NK-EVs curb CCL4-induced liver fibrosis in mice by inhibiting TGF-β1-induced hepatic stellate cells activation (12). Also miR-207-containing NK-EVs alleviate depression-like symptoms in mice (13).

Interestingly, recent studies have identified the role of biologically active NK-EV miRNAs, such as miR-186, which impaired neuroblastoma tumour growth and inhibited immune escape mechanisms by targeting the TGF-β pathway (14); or miR-3607-3p, which inhibited pancreatic cancer, presumably by targeting its putative target IL-26 (15). These reports suggest the modulatory role of NK-EV-miRNAs to inhibit cancer progression. However, little is known regarding the overall small RNA composition of human NK-EVs, and their impact on the regulation of immune responses remains far from being fully elucidated.

In this study, we have set up a model to study human primary NK-EVs. We have analysed the small RNA content of resting and *in vitro* activated NK cells and their secreted EVs by next-generation sequencing (NGS). We show that NK-EVs have specific miRNA repertoire and post-transcriptional modification (PtM) patterns, which differ from that of their parental cells. Analyses of the NK-EV miRNA signature revealed an enrichment of Th1 function-related miRNAs that target key T cell mRNA molecules. One example is GATA-3, as its downregulation leads to T-bet de-repression. Other identified miRNAs are involved in DC presentation and co-stimulatory functions. We further showed that NK-EVs promote T cell activation followed by IFN-γ and IL-2 production and induced DC expression of MHC-II and CD86 in DCs. Finally, *in vivo* nanoparticle-based delivery of the NK-EV enriched miRNAs miR-10b-5p, miR-92a-3p and miR-155-5p partially reproduces the effects of NK-EV treatment in T cell responses, suggesting an involvement of these miRNAs in effector functions. These findings shed light on the antitumor effect of NK-EVs and support their use as therapeutic tools.

## RESULTS

### NK-derived EVs bear a specific miRNA signature different from their secreting cells

To identify the miRNA repertoire of resting- and *in vitro*-activated NK cells and their load into EVs, we set up the protocol shown in **Figure 1—figure supplement 1A**. Briefly, NK cells were enriched from isolated peripheral blood mononuclear cells (PBMCs) of human healthy dono’s buffy coats by adding a mixture of IL-12 and IL-18 together with irradiated feeder cells. Six days after, NK cell expansion was tested by flow cytometry, and cells were kept in culture with IL-2 for a further 48 h, in the absence of feeder cells, before NK cell isolation **(Figure 1—figure supplement 1A)**. EVs accumulated for 72 h from 15×10^6^ NK cells were purified by serial centrifugation as previously described (16) and tracked (EV-TRACK ID: EV210234). RNA was isolated from resting NK cells (directly isolated without being cultured), *in vitro* activated NK cells and small EVs released from activated NK cells **(Figure 1—figure supplement 1A,C)**. Vesicles isolated from activated NK cells were characterized using Nanosight showing a mode size ranging from 165 to 209 nm among donors **(Figure 1—figure supplement 1D-F)**. Also, biochemical analyses showed the NK-EV expression of the exosomal markers CD63 and Tsg101 and the absence of the EV-excluded marker Calnexin **(Figure 1—figure supplement 1G)**.

Small RNA sequencing analyses showed that resting and activated cells display a distinct miRNA profile that differs to that of cultured NK cells-secreted EVs, as shown in Heatmap and PC plots **(Figure 1A—figure supplement 1H and Supplementary Table 1)**. A total of 130 miRNAs were differentially expressed between activated NK cells and NK-EV released miRNAs; 96 miRNAs being enriched in NK-EVs and 34 miRNAs being more abundant in NK cells than in NK-EVs **(Figure 1B)**. Such differential expression suggested the existence of mechanisms of specific miRNAs sorting into EVs derived from NK cells, in agreement with published data (17–21). Also, the repertoire found in resting NK cells was different from that of *in vitro* activated NK cells **(Figure 1A—figure supplement 1I and Supplementary Table 1).** A total of 70 miRNAs were upregulated upon activation, while 98 miRNAs decreased their levels after cytokine stimulation **(Figure 1A—figure supplement 1I)**. These data are in agreement with extensive miRNA remodelling occurring upon T lymphocyte activation (22).

**Fig. 1.**
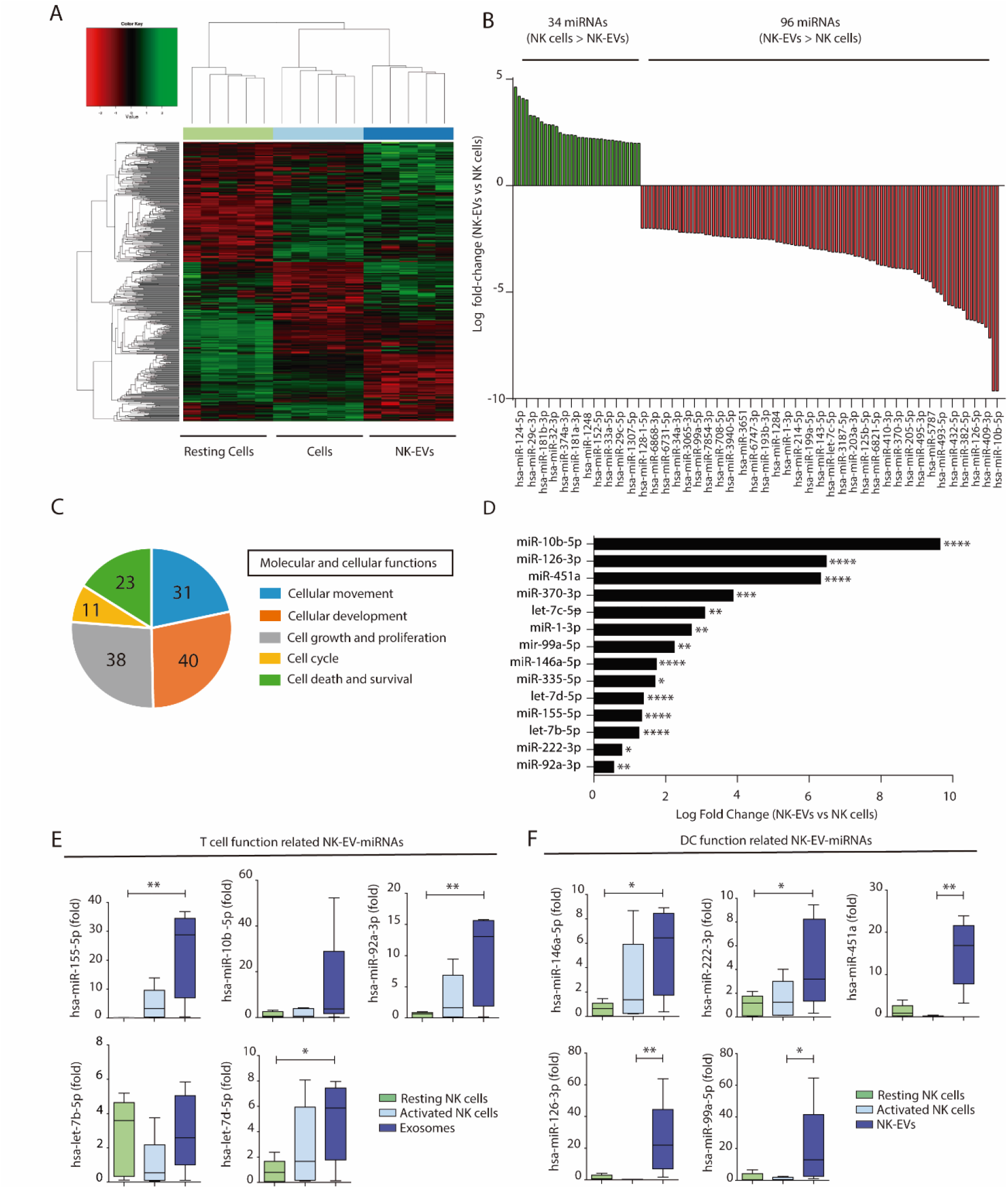
NK-EVs have a specific miRNA signature and are enriched in miRNAs related with Th1 functions. **(A)** Heat map showing small RNA sequencing analysis of miRNAs differentially expressed in resting and activated human NK cells and their secreted EVs. Data are from NK cells isolated from five healthy donors. Significance was assessed using the Benjamini-Hochberg procedure and only miRNAs with an adjusted p-value < 0.05 are shown. (**B**) Histogram plot showing logarithmic fold-increase expression between activated NK cells and their released EVs. Only fold-changes with adjusted P-value < 0.05 and log fold-change > 2 are represented. MiRNAs significantly more expressed in cells than in their secreted exosomes are shown in green, while miRNAs significantly more represented in the EV fraction are shown in red. (**C**) Summary of molecular and cellular functions targeted by NK-EV miRNAs identified by unbiased Ingenuity Pathway Analysis (IPA). The numbers indicate the molecules targeted by miRNAs over-represented in NK-EVs compared to NK cells. (**D**) Log fold-change in the small RNAseq expression of miRNAs significantly overexpressed in NK-EVs compared to their secreting cells, related to Th1 functions. Significance was assessed using Benjamini-Hochberg adjusted p-values; *P<0.05,**P<0.01, ***P<0.001, ****P<0.0001. (**E,F**) Quantitative real-time PCR of NK-EV upregulated miRNAs in resting NK cells, activated NK cells and secreted EVs. Bars represent the mean ± SEM of cells and vesicles obtained from 5 healthy donors, normalized to the small nucleolar RNU5G. Data show the validation of CD4+ T cell (**E**) and DC (**F**) function related miRNAs, obtained by the 2-ΔΔCt method, using Biogazelle software. Significance was assessed by One-Way ANOVA Dunńs test; *P<0.05, **P<0.01.

Unbiased analysis of the pathways potentially affected by miRNAs shuttled into NK-EVs using the Ingenuity Pathway Analysis revealed that the putative mRNA targets of NK-EV enriched miRNAs were involved in cellular development and movement, cell growth and proliferation, cell death, survival and cell cycle **(Figure 1C)**. *In silico* mRNA target analyses for NK-EV-miRNAs identified putative target molecules related to immune signalling and Th1 responses, targeting CD4^+^ T lymphocytes and DCs among other immune effectors **(Supplementary Table 2)**. Selected miRNAs identified in this screening **(Figure 1D)** were validated by qPCR **(Figure 1E,F)**.

### Post-transcriptional modifications in NK cells and EV miRNAs

Since changes in the miRNA repertoire were linked to post-transcriptional modifications (PtMs) of miRNAs during T lymphocyte activation (22), we analysed the global PtMs signatures of miRNAs from resting NK cells, activated NK cells and released NK-EVs **(Figure 2A-C)**. These analyses revealed a complex pattern of miRNA PtMs in human NK cells and released small EVs. Most PtMs appeared at the edges of the canonical sequence, particularly at the terminal nucleotide (position 0) and the two flanking nucleotide positions (−1 and +1). The 3p-end of miRNAs was significantly more prone to PtMs **(Figure 2A)** than the 5p-end in every sample **(Figure 2B—figure supplement 2A)**. Also, non-templated mono-additions of nucleotides were much more abundant than poly-additions **(Figure 2D—figure supplement 2B)**. Thus, we focused our analyses on the 3p-end miRNÁs PtMs, in particular single nucleotide additions accumulated in positions −1 to +1. Interestingly, miRNAs from the different fractions (resting NK cells, activated NK cells and NK-EVs) exhibited clear PtM signature differences **(Figure 2C,D).** Cytosine addition was the most common modification, followed by adenine terminal insertion. Upon NK cell activation, miRNAs displayed reduced adenylation (positions −1, +1) and cytosylation (positions −1, +1). Comparing activated NK cells with the NK-EVs they release, we detected increased levels of cytosylated miRNA reads (positions 0, +1), and a decrease in guanosine additions (positions −1, 0). Although other significant changes in miRNA PtMs are observed in the different groups under study, they are supported by fewer number of reads.

**Fig. 2.**
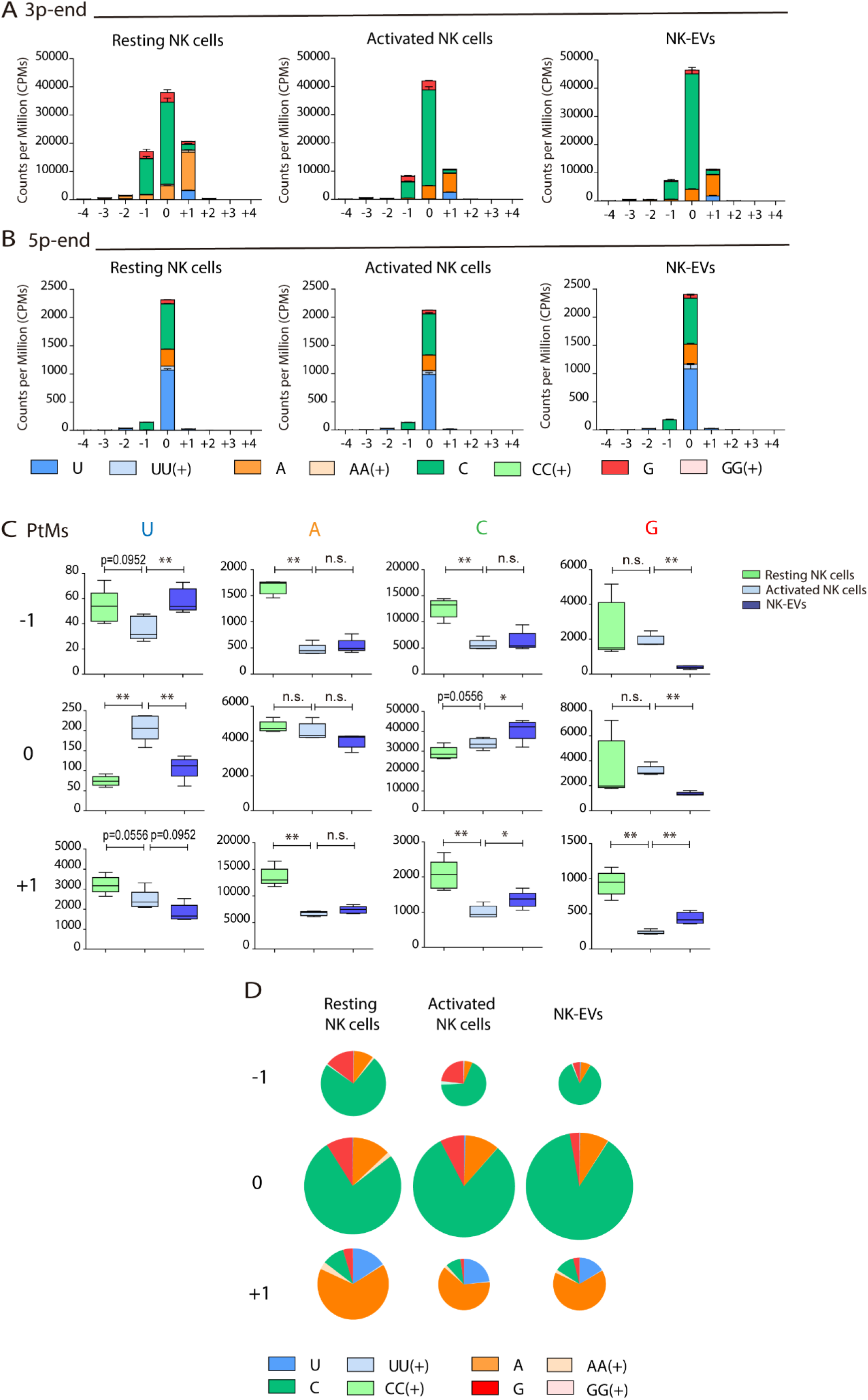
Resting NK cells, activated NK cells and their released EVs contain miRNAs with a different PtM signature. (**A,B**) Bar charts showing the PtM profile of miRNAs from resting NK cells, activated NK cells and NK-EVs, expressed in normalized Counts per Million (CPMs) at the 3p-end (**A**) and 5p-end (**B**), respectively. Modifications from the canonical sequence in positions ranging from −4 to +4 are represented. (**C**) Box and whiskers plots show the additions of U, A, C and G at the indicated positions (ranging from −1 to +1) in miRNAs from resting NK cells, activated NK cells and NK-EVs. Significance was assessed, comparing resting and activated NK cells and activated NK cells with their released small EVs using non-parametric t-test; *P<0.05, **P<0.01. (**D**) Pie charts show the proportion of the different modifications (U, A, C and G) for each position (from −1 to +1) for resting NK cells, activated NK cells and NK-EV miRNAs, respectively. The area of each pie chart is proportional to the total number of reads bearing the specified PtMs at the indicated positions.

Given that the number of detected PtMs are influenced by the level of expression of individual miRNAs, we next studied the fraction of reads with PtMs within individual miRNAs. Remarkably, analysing the percentage of reads with PtMs for the miRNAs specifically enriched in NK-EVs **(Figure 3A,** right panel), we found a significantly higher proportion of PtMs from these miRNAs in NK-EVs than in their secreting-activated NK cells. This suggests that miRNA molecules with PtMs are more likely to be included into EVs. In fact, 5 out of 6 of the most enriched NK-EV miRNAs show a higher number of reads with PtMs in NK-EVs compared to activated NK cells **(Figure 3B)**. We next analysed the specific nucleotide mono-additions in miRNAs preferentially targeted into NK-EVs **(Figure 3C)**. We found that both adenylation and cytosylation of canonical miRNA sequences accounted for the observed accumulation of PtMs in the NK-EV fraction. The levels of guanosylated miRNAs were very low in all samples, although more represented in resting NK cells, while no significant differences were observed in the levels of uridine additions, among the samples analysed.

**Fig. 3.**
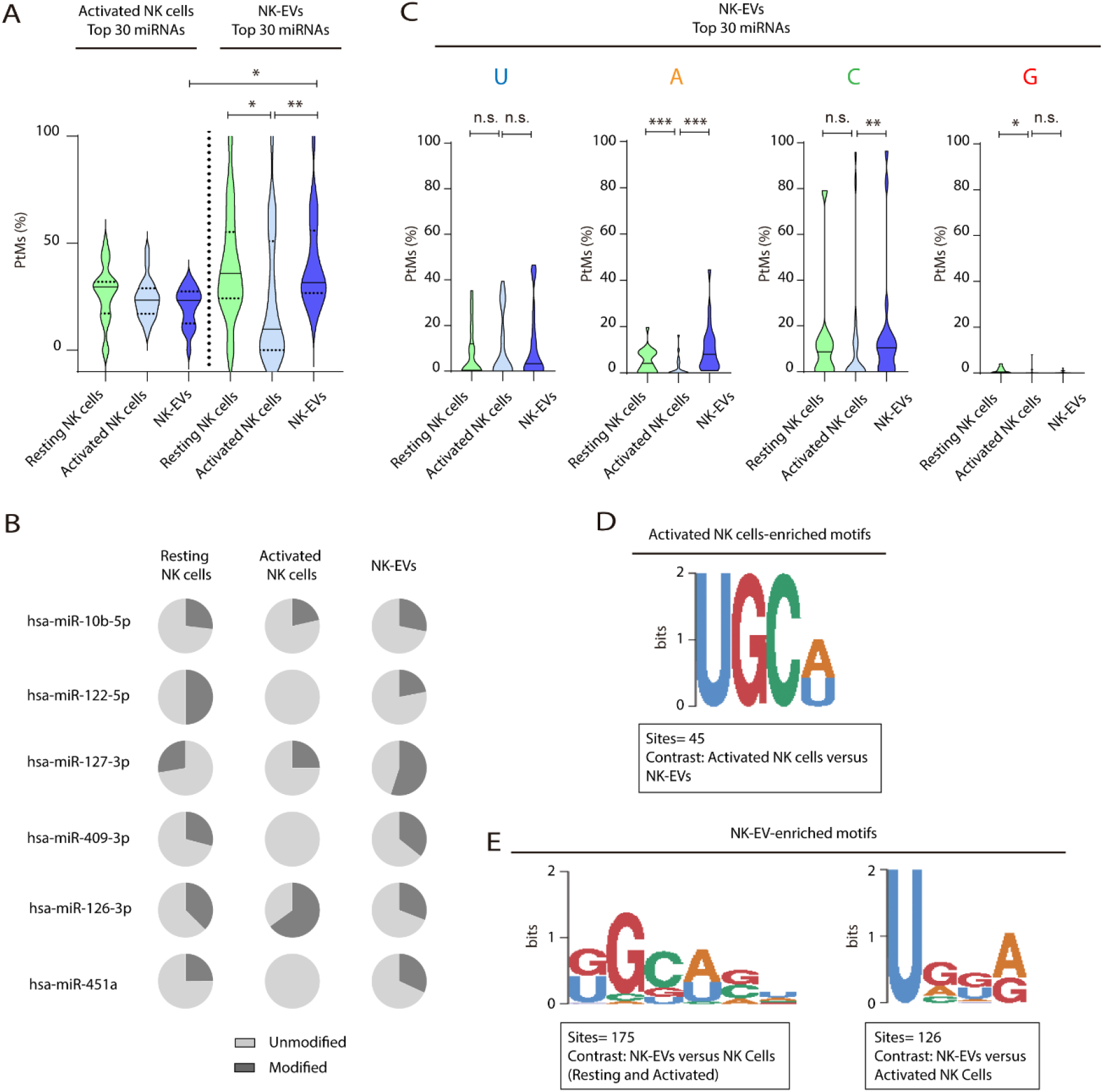
NK-EV miRNAs are enriched in non-templated nucleotide additions and specific short motifs. **(A)** Violin plots represent the fraction of miRNA reads with PtMs. Analysis was performed comparing the most differentially expressed miRNAs in activated NK cells compared to NK-EVs. The left panel summarizes the data obtained from the 30 miRNAs more expressed in the activated NK cell fraction, while the right panel shows the percentage of reads with PtMs from the 30 miRNAs with a higher expression in NK-EVs compared to their secreting activated NK cells. Individual miRNAs with less than 10 reads were excluded from the analysis. The median value is shown as a solid line and quartiles represented as dotted lines. Significance was assessed with the Kruskal-Wallis test; *P<0.05, **P<0.01. (**B**) Pie charts showing the proportion of the modified (dark gray) and unmodified (light gray) fractions respectively, for the six miRNAs more differentially enriched in activated NK-derived EVs compared to their secreting cells. (**C**) Violin plots represent the fraction of miRNA reads with U, A, C or G mono-additions at positions −1, 0 and +1. Analysis was performed for the 30 miRNAs more enriched in NK-EVs, comparing the PtM signature in resting cells and activated cells, and activated cells and their released NK-EVs. Significance was assessed with the non-parametric t-test Wilcoxon matched-pairs signed ranked test; *P<0.05, **P<0.01,***P<0.001. (**D,E**) Over-represented motifs in activated NK cells (**D**) and NK-EV-miRNAs (**E**), using the ZOOPS model. For each data set, all miRNAs annotated in miRBase 21 were used as background. Short motifs with adjusted E-value<0.2 are shown.

Altogether, accumulation of PtMs in miRNAs enriched in NK-EVs suggests an active PtMs-dependent mechanism of specific miRNA sorting into EVs. However, the accumulation of modified miRNAs in NK-EVs was not observed in the case of the most expressed NK-EV miRNAs **(Figure 3—figure supplement 3A)**, indicating additional layers of complexity in NK-EV-miRNA specific packaging, e.g. passive sorting of the most abundant miRNAs.

Specific short sequence miRNA motifs are over-represented in EVs. Binding of these motifs to RNA binding proteins, e.g. heterogeneous nuclear ribonucleoprotein A2/B1 (hnRNPA2B1), promotes miRNA sorting into EVs (18). Bioinformatics sequence analyses showed over-represented short motifs in resting NK cells (UGCUG, **Figure 3—figure supplement 3B**), activated NK cells (UGCA, **Figure 3D**), and released NK-EVs (GGCAGU and UGGA, **Figure 3E**), respectively. Interestingly, the short EV-associated motif identified in T cell-derived small EVs GGAG was over-represented in NK-EV-miRNAs, and the tetra-nucleotide motif UGCA was also identified as a “cell retention motif” in T cells, pointing towards similar mechanisms of miRNA sorting in lymphocytes (23).

Our analyses of NK-EV enriched specific miRNAs suggest that specific miRNAs packaging into NK-EVs may involve different mechanisms, including non-templated nucleotide additions to miRNA canonical sequences and short-motif miRNA recognition by RNA-binding proteins. Active targeting of specific miRNAs for EV release implies a putative role for these NK-EV-miRNAs in intercellular communication and regulatory functions.

### NK-EVs promote Th1-like responses with increased T-bet expression and IFN-γ and IL-2 release

To dissect the putative immunomodulatory effects of shuttled NK-EV miRNAs, analyses of the individual miRNAs, either more abundant in the NK-EV fraction **(Figure 3—figure supplement 3C)** and more differentially enriched in NK-EVs as compared to activated NK producing cells, **(Figure 3—figure supplement 3D),** and their *in silico*-predicted mRNA targets **(Supplementary Table 2)** were carried out. As previously pointed out, miRNAs with key roles in immune function regulation, in particular with T cell responses, such as miR-10b-5p, miR-155-5p or miR-92a-3p were highly expressed in NK-EVs.

To unravel the effects of NK-EVs on T cell function, CD4^+^ T cells were isolated from human buffy coats and cultured either in the presence or absence of purified NK-EVs. An initial profiling of the cytokines secreted by NK-EV treated CD4^+^ T cells in non-cytokine polarizing conditions showed high levels of Th1-function related cytokines, including IFN-γ and IL-2 **(Figure 4—figure supplement 4A)**. Unbiased IPA analysis had identified cell death and survival as pathways potentially affected by NK-EV enriched miRNAs, hence we evaluated CD4^+^ T cell survival **(Figure 4—figure supplement 4B,C)**. We found no significant differences in overall CD4^+^ T cell survival after NK-EV treatment.

Further *in vitro* studies were performed to address the effects of NK-EV addition in non-polarizing and Th1-skewing conditions using a mixture of IL-2 and IL-12. Cultured CD4^+^ T cells were analysed after three and six days, respectively. Flow cytometry analysis of cultured CD4^+^ T cells showed an increase in the percentage of IFN-γ producing T cells in the presence of NK-EVs in non-polarizing conditions. However, the effects were not evident in Th1-polarizing conditions **(Figure 4A)**. Sandwich-ELISA analysis of cultured cells supernatants revealed an increase in the levels of soluble IFN-γ in NK-EV treated T cells in a non-cytokine polarizing milieu **(Figure 4B).** In addition, incubation with NK-EVs also induced T-bet expression in non-cytokine polarized T cells **(Figure 4C,D—figure supplement 4D),** correlating with a downregulation of the NK-EV miRNA target Gata-3 mRNA levels at day 3, which acts as a T-bet suppressor **(Figure 4E)** (24). The increase of the NK-EV identified miRNAs miR-10b-5p and miR-92a-3p, which have crucial functions in T cell responses **(Supplementary Table 2)** was confirmed by RT-qPCR in non-polarized culture conditions but not in cytokine Th1-polarizing cultures, after 3 days of incubation with NK-EVs **(Figure 4F)**. Altogether, NK-EV miRNA driven Gata-3 downregulation may promote T-bet de-repression and subsequent Th1 reprogramming in T cells.

**Fig. 4.**
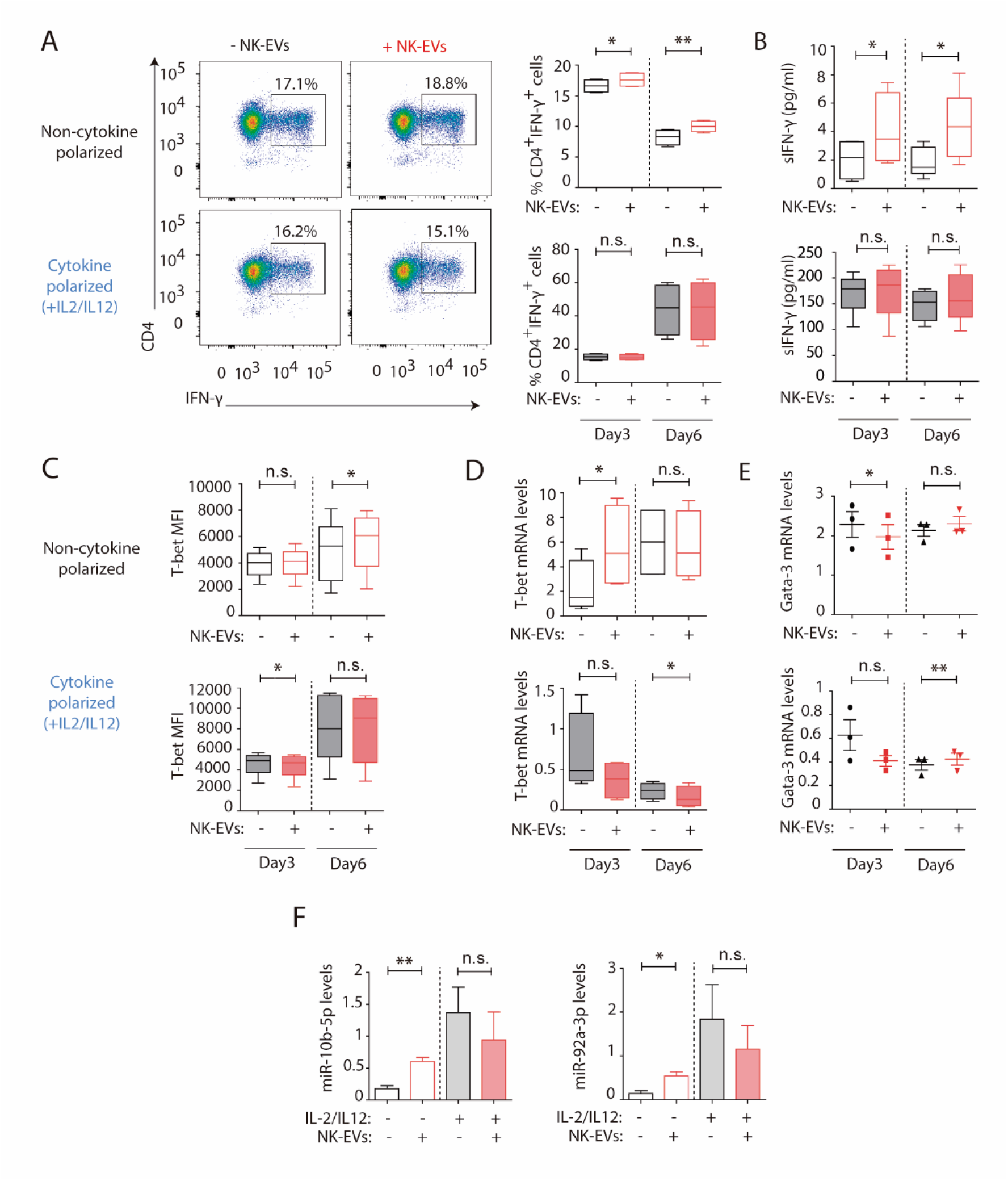
NK-EVs promote Th1 differentiation via Gata-3 downmodulation and T-bet de-repression correlating with increased levels of miR-10b-5p and miR-92a-3p. CD4+ T cells isolated from healthy human buffy coats were cultured either in non-polarizing or in cytokine Th1-polarizing conditions, with a mixture of IL-2 and IL-12, in the presence or absence of NK-EVs. (**A**) Flow cytometry analysis of isolated CD4+ T lymphocytes incubated under non-cytokine polarizing (upper panels) and Th1 cytokine-polarizing (with a mixture of IL-2 and IL-12, lower panels) conditions. A representative experiment is shown. Box and whiskers plots show the expression of CD4 and IFN-γ in gated live cells (min to max and median values), after addition of NK-EVs. Right panel plots show the quantification of n ≥ 4 independent experiments. Significance was assessed with Paired Student’s t-test; *P<0.05, **P<0.01. (**B**) ELISA quantification of soluble IFN-γ in supernatants from cultured cells in the indicated conditions (unpolarized, upper panel; cytokine polarized, lower panel). The chart shows the median concentration from n ≥ 4 independent experiments. Significance was assessed with Paired Student’s t-test; *P<0.05. (**C**) Flow cytometry analysis of isolated CD4+ T cells cultured in the different conditions. Graphs show the Mean Fluorescence Intensity of T-bet in gated live single CD4+ T cells after 3 and 6 days of culture respectively. Significance was assessed with Paired Student’s t-test; *P<0.05. (**D,E**) Quantitative real-time PCR at days 3 and 6 showing mRNA levels of T-bet (**D**) and Gata-3 (**E**), respectively, normalized to GAPDH and β-actin. Significance was assessed by Paired Student’s t test; *P<0.05. (f) Quantitative real-time PCR at day 3 to detect miRNA levels in CD4+ T cells after culture in the indicated conditions. MiR-10b-5p (left panel) and miR-92a-3p (right panel) relative expression is shown, and normalized to RNU1A1 and RNU5G. Significance was assessed by Paired Student’s t test; *P<0.05.

The activation of CD4^+^ T cells by NK-EVs was also evaluated. NK-EVs promoted T cell activation, as revealed by increased CD25 expression in non-polarizing conditions; whereas these effects were not further increased in a Th1-cytokine polarized milieu **(Figure 5A)**. Increased IL-2 release upon incubation with NK-EVs was confirmed by ELISA, 3 days after culture **(Figure 5B).** Since CD25 is also a hallmark of regulatory T cells (Treg), we next analysed the expression of Foxp3 and IL-10 production in NK-EV treated T lymphocytes **(Figure 5C).** Although no significant differences were found, increased Foxp3 levels (p-value < 0.1) in non-polarized NK-EV treated cells were observed compared to non-EV treated cells. However, IL-10 secretion was not affected by NK-EV treatment, and was even reduced 3 days after culture in polarizing conditions. These data suggest that additional levels of regulation participate in T cell responses, and that NK-EVs may limit suppressive Treg functions in Th1 skewing conditions.

**Fig. 5.**
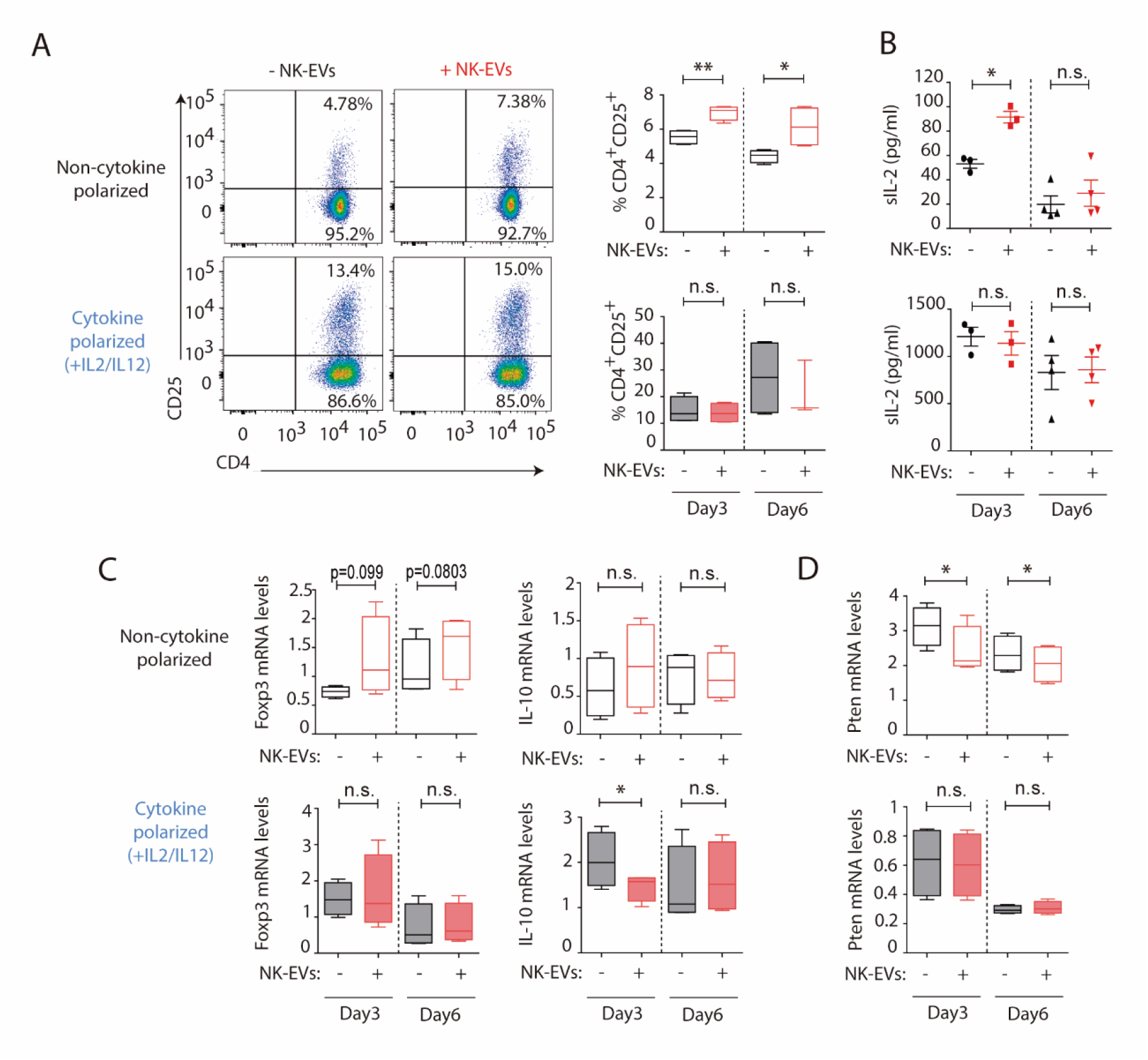
NK-EVs promote CD4+ T cell activation and IL-2 release but not Treg responses. (**A**) Flow cytometry analysis of isolated CD4+ T cells incubated under non-cytokine polarizing (upper panel) and Th1 cytokine-polarizing (with a mixture of IL-2 and IL-12, lower panel) conditions. Dot plots show the expression of CD4 and CD25 in gated single live cells ± SEM, after addition of NK-EVs. Plots show the quantification of n ≥ 4 independent experiments. Significance was assessed with Paired Student’s t-test; *P<0.05, **P<0.01. (**B**) ELISA quantification of soluble IL-2 in supernatants from CD4+ cultured T cells in the indicated conditions (unpolarized, upper panel; cytokine polarized, lower panel). The graph shows the mean concentration from n ≥ 3 independent experiments. Significance was assessed by Paired Student’s t-test; *P<0.05. (**C,D**) Quantitative real-time PCR showing FoxP3 (**C**, left); IL-10 (**D**, right) and Pten (**D**) mRNA levels at days 3 and 6 in CD4+ T cells after culture in the indicated conditions. Relative expression is shown, normalized to GAPDH and β-actin. Significance was assessed by Paired Student’s t test; *P<0.05.

Analysis of additional putative NK-EV miRNA targets described in **Supplementary Table 2** was carried out **(Figure 5D—figure supplement 5)**. Functional regulatory molecules were downregulated by NK-EVs, including PTEN **(Figure 5D)**, which is involved in T cell homeostasis; IL-6, which was reduced after NK-EV incubation (p-value < 0.1)); TGF-β and Ship1 **(Figure 5—figure supplement 5A-F)**. On the contrary, the pleiotropic TNF-α cytokine was upregulated shortly after NK-EV incubation in non-polarizing conditions, but decreased in Th1 cytokine-polarizing conditions **(Figure 5—figure supplement 5E).** Altogether our data indicate a complex regulation of T cells responses mediated by NK-EVs.

### NK-EVs promote DC-mediated Th1 responses, enhancing monocytes and moDC presentation and costimulatory functions

Dendritic cells are critical for the establishment of effective T-cell responses (25). To ascertain the role of NK-EVs in DC function, human monocytes were obtained, and cultured either in DC polarizing conditions (supplemented with a mixture of GM-CSF and IL-4) or without cytokines (26). The effects of the addition of NK-EVs were assessed in both experimental conditions (monocytes and monocyte-derived DCs, moDCs). Flow cytometry analyses revealed that, although no significant differences were observed in cell survival and apoptosis **(Figure 6—figure supplement 6A,B)**, the presentation and co-stimulatory capacities of DCs were enhanced by NK-EVs exposure **(Figure 6A-D).** Indeed, both NK-EV-treated monocytes and moDCs showed increased levels of MHC-II **(Figure 6A)** and the co-stimulatory CD86 molecule **(Figure 6B)**. Additionally, non-cytokine polarized monocytes exhibited a higher DC-polarization, marked by an increased percentage of CD11c^+^DC-Sign^+^ cells after incubation with NK-EVs **(Figure 6C)**. Interestingly, NK-EVs addition correlated with increased levels of IFN-γ and IL-12 secretion in monocytes, but not in moDCs that exhibited very low levels of these cytokines **(Figure 6D,E)**.

**Fig. 6.**
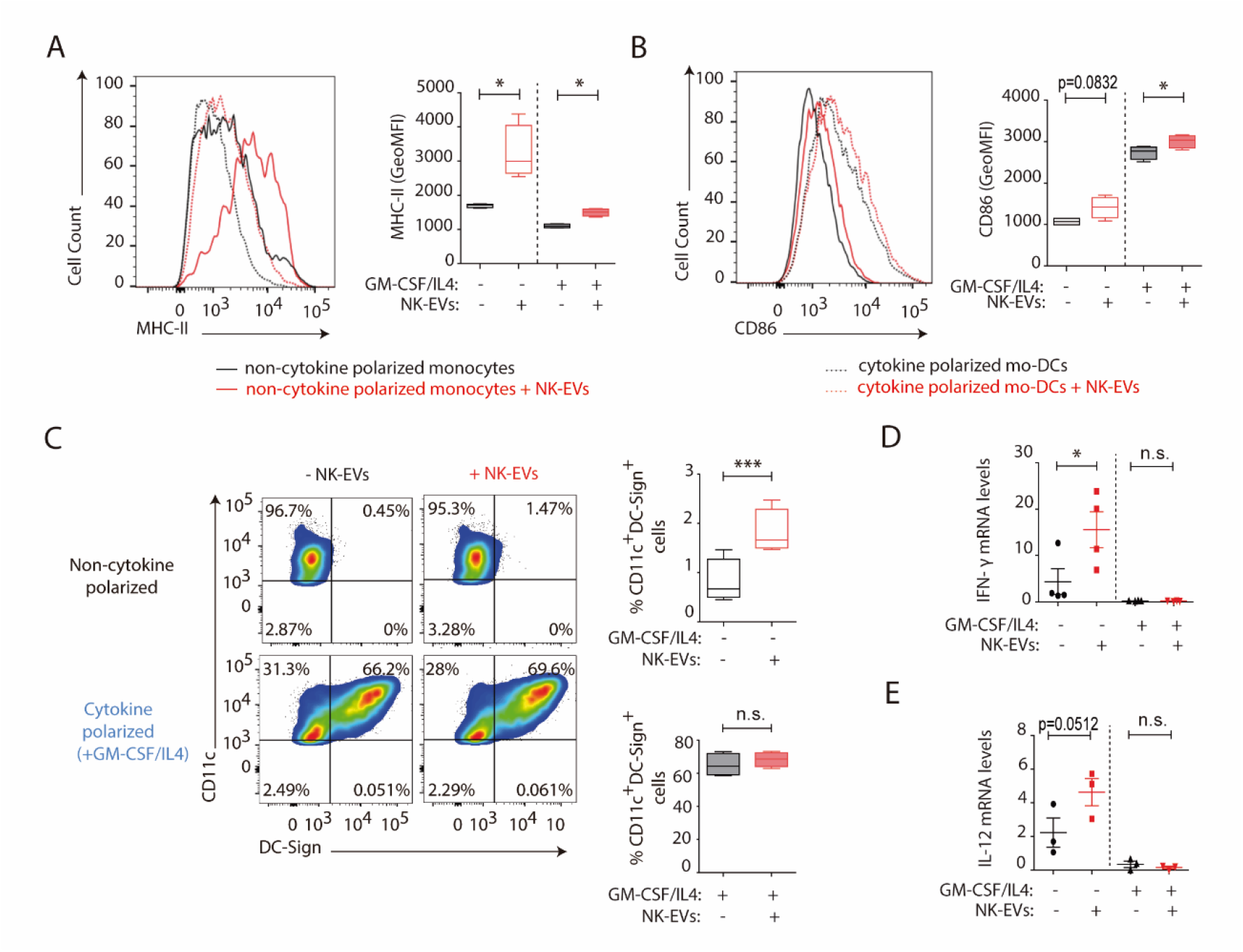
NK-EVs enhance monocyte and moDCs polarization, costimulatory and presentation activities. CD14+ monocytes isolated from buffy coats from healthy donors were cultured under non-polarizing and cytokine monocyte-derived DC-polarizing conditions, with a mixture of GM-CSF and IL-4, with or without NK-EVs. (**A,B**) Flow cytometry analysis of isolated CD14+ monocytes in the different culture conditions. Histogram plots show the expression of MHC-II (**A**) and CD86 (**B**), after 6 days of culture in the indicated conditions, either in the presence (dotted line) or absence (solid line) of polarizing cytokines. A representative plot of n ≥ 4 experiments of the Mean Fluorescence Intensity (MFI) of these proteins in live cultured cells (left panel), and their quantification (right panel) are shown. Significance was assessed by Paired Student’s t test; *P<0.05. (**C**) Flow cytometry analysis of isolated CD14+ monocytes in the different culture conditions. Dot plots show the expression of CD11c and DC-Sign in the indicated culture conditions (left panel) and their quantification (right panel). Significance was assessed by Paired Student’s t test; ***P<0.001. (**D,E**) Quantitative real-time PCR showing IFN-γ (**D**) and IL-12 (**E**) mRNA levels in CD14+ cells 6 days after culture in the indicated conditions. Relative expression is shown, normalized to GAPDH and β-actin. Significance was assessed by Paired Student’s t test; *P<0.05.

Several putative targets for NK-EV miRNAs with important functions in DCs **(Supplementary Table 2)** were analysed after treatment with NK-EVs, including KLF13, Foxo1, Notch1 and VEGFR2, but no significant expression differences were found **(Figure 6—figure supplement 6C-F).**

### NK-EV miRNAs partially recapitulate NK-EV polarizing functions

To address whether miRNAs contained in NK-EVs may account for the observed Th1 immune deviation, nanoparticles bearing either individual miRNAs or a combination of several miRNA molecules were synthesized. The effects of *in vivo* footpad administration of nanoparticles carrying the NK-EV-miRNAs miR-10b-5p, miR-92a-3p and miR-155-5p were analysed. Interestingly, splenocytes isolated from nanoparticle-treated mice activated *in vitro* with anti-CD3 and anti-CD28 antibodies showed increased T cell activation after miRNA-bearing treatment compared to control nanoparticles, assessed by increased CD25 expression in CD4^+^ lymphocytes **(Figure 7A,B)**. qPCR analysis of total splenocytes **(Figure 7C)** and popliteal draining lymph nodes **(Figure 7D)** showed increased mRNA levels of the Th1 hallmark IFN-γ after footpad administration of NK-EV miRNAs, in particular with miR-155 and a mixture of all three NK-EV miRNAs. Increased secretion of IFN-γ was also confirmed in supernatants from *in vitro* activated splenocytes by sandwich-ELISA **(Figure 7E)**. Although no significant differences were found in the percentage of IFN-γ expressing CD4^+^ T cells **(Figure 7F, lower panel)**, we could observe a similar tendency in T cells from mice treated with miR-10b-5p and miR-92a-3p, that exhibited a significantly enhanced expression (Mean fluorescence Intensity) of this cytokine upon TCR-stimulation, compared to non-miRNA treated cells **(Figure 7F, upper panel).** The effects of miR-10b-5p, miR-92a-3p and miR-155-5p delivery in CD8^+^ T cells IFN-γ secretion of this cytokine were more robust **(Figure 7G).** The levels of T-bet were slightly increased in miRNA-treated mice **(Figure 7—figure supplement 7A)**, although the differences were not significant, likely due to the kinetics of T-bet upregulation after Th1 commitment (27). Importantly, as observed in the experiments after NK-EV treatment *in vitro*, no differences were observed in the Treg compartment upon nanoparticle-based miRNA delivery, as addressed by the percentage of CD4^+^CD25^+^Foxp3^+^ cells **(Figure 7—figure supplement 7B)** and the levels of IL-10, neither in the spleen **(Figure 7—figure supplement 7C)**, nor in the popliteal draining lymph nodes **(Figure 7—figure supplement 7D)**. ELISA analysis of supernatants splenocytes from treated mice showed no significant differences in the levels of IL-2 secretion **(Figure 7—figure supplement 7E)**. Low levels of both IFN-γ and IL-2 cytokines were detected in serum from treated mice, however miR-10b-5p and miR-155-5p bearing nanoparticles promoted a slight increase of these cytokines **(Figure 7—figure supplement 7F,G)**.

**Fig. 7.**
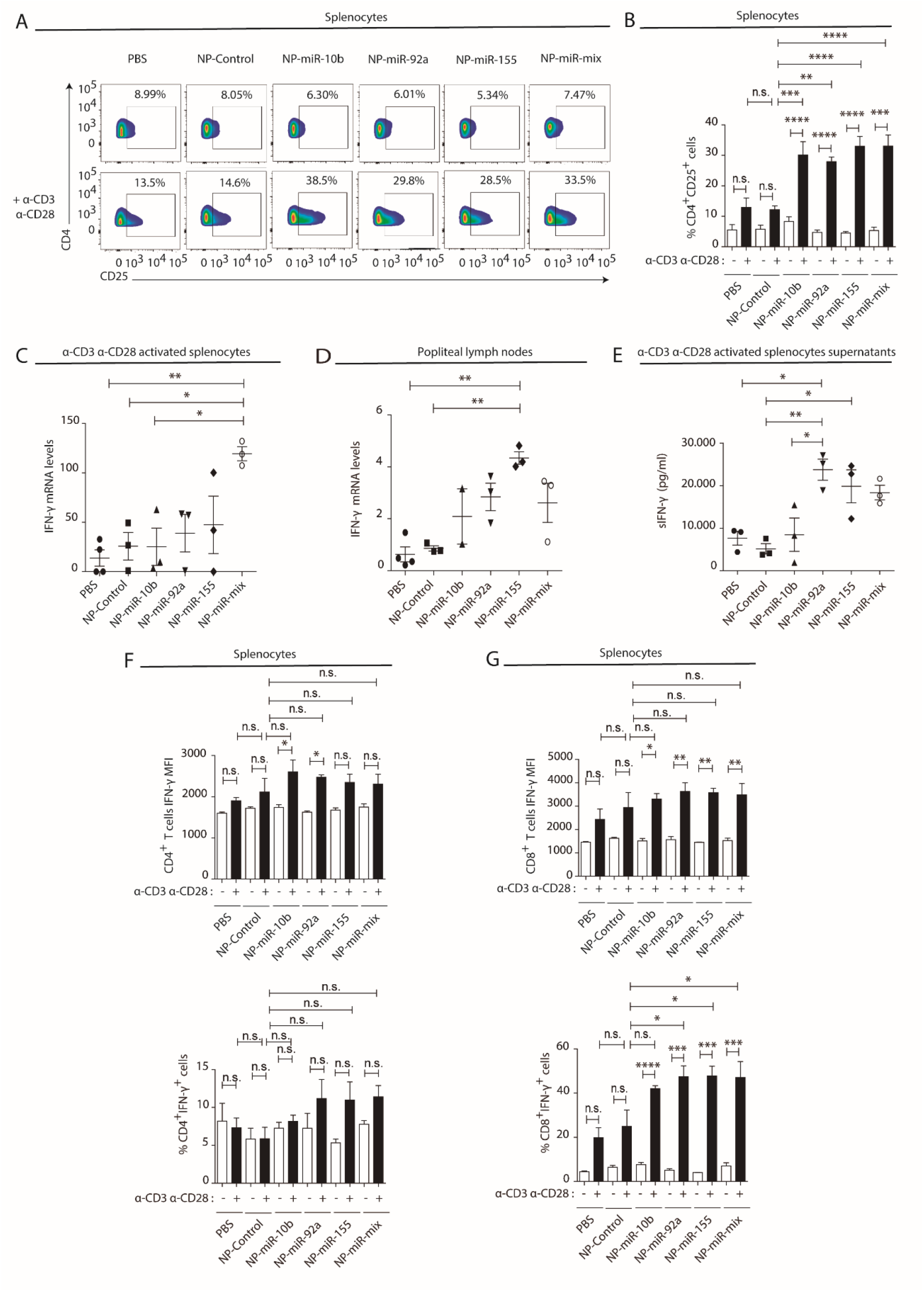
NK-EV T-cell function related miRNAs partially mimic NK-EV mediated effects *in vivo*. NPs bearing the identified NK-EV miRNAs were generated and their functional effects were evaluated in vivo after footpad injection of control and miRNA-loaded NPs in wild-type C57/BL/6 mice. (**A**) Flow cytometry analysis of live splenocytes isolated 6 days after footpad injection with the indicated control or miRNA-loaded NPs. Isolated cells were either left unstimulated (upper panel) or incubated for 16 h with anti-CD3 and anti-CD28 antibodies (lower panel) before analysis. Density plots show the expression of CD4 and CD25 in TCRβ+ CD4+ gated live T lymphocytes. A representative plot of n ≥ 3 mice is shown. (**B**) Bars summarize the quantification of the percentage of CD4+ CD25+ T cells shown in (**A**) from n ≥ 3 mice in untreated splenocytes (white) and anti-CD3 plus anti-CD28 stimulated splenocytes (black). Significance was assessed by Two-way ANOVA, followed by Tukeýs test; **P<0.01, ***P<0.001, ****P<0.0001. (**C,D**) Quantitative real-time PCR showing IFN-γ mRNA levels in anti-CD3 and anti-CD28 activated splenocytes (**C**) and draining popliteus lymph nodes (**D**), 6 days after footpad nanoparticle-based miRNA delivery. Relative expression is shown, normalized to GAPDH and β-actin. Significance was assessed by One-way ANOVA Bonferroni test; *P<0.05, **P<0.01. (**E**) ELISA quantification of soluble IFN-γ in supernatants from splenocytes harvested and isolated 6 days after miRNA delivery and cultured with anti-CD3 and anti-CD28 antibodies for 16 h. The graph shows the mean concentration from n ≥ 3 independent experiments. Significance was assessed by One-way ANOVA Bonferroni test; *P<0.05, **P<0.01. (**F,G**). Upper panel bar charts show the Mean Fluorescence Intensity (MFI) expression of IFN-γ and lower panels the percentage of IFN-γ expressing cells, analysed by flow cytometry in CD4+ T cells (**F**), CD8+ T cells (**G**). Splenocytes were either left unstimulated (white) or stimulated with anti-CD3 plus anti-CD28 antibodies for 16 h (black). Significance was assessed by Two-way ANOVA, followed by Tukeýs test; *P<0.05,**P<0.01, ***P<0.001, ****P<0.0001.

In summary, activated NK cells release a specific set of EV-miRNAs, likely by mechanisms involving PtMs and specific-short miRNA motifs recognition, which are related with Th cells effector functions and may therefore contribute to orchestrate the immune response **(Figure 8)**.

**Fig. 8.**
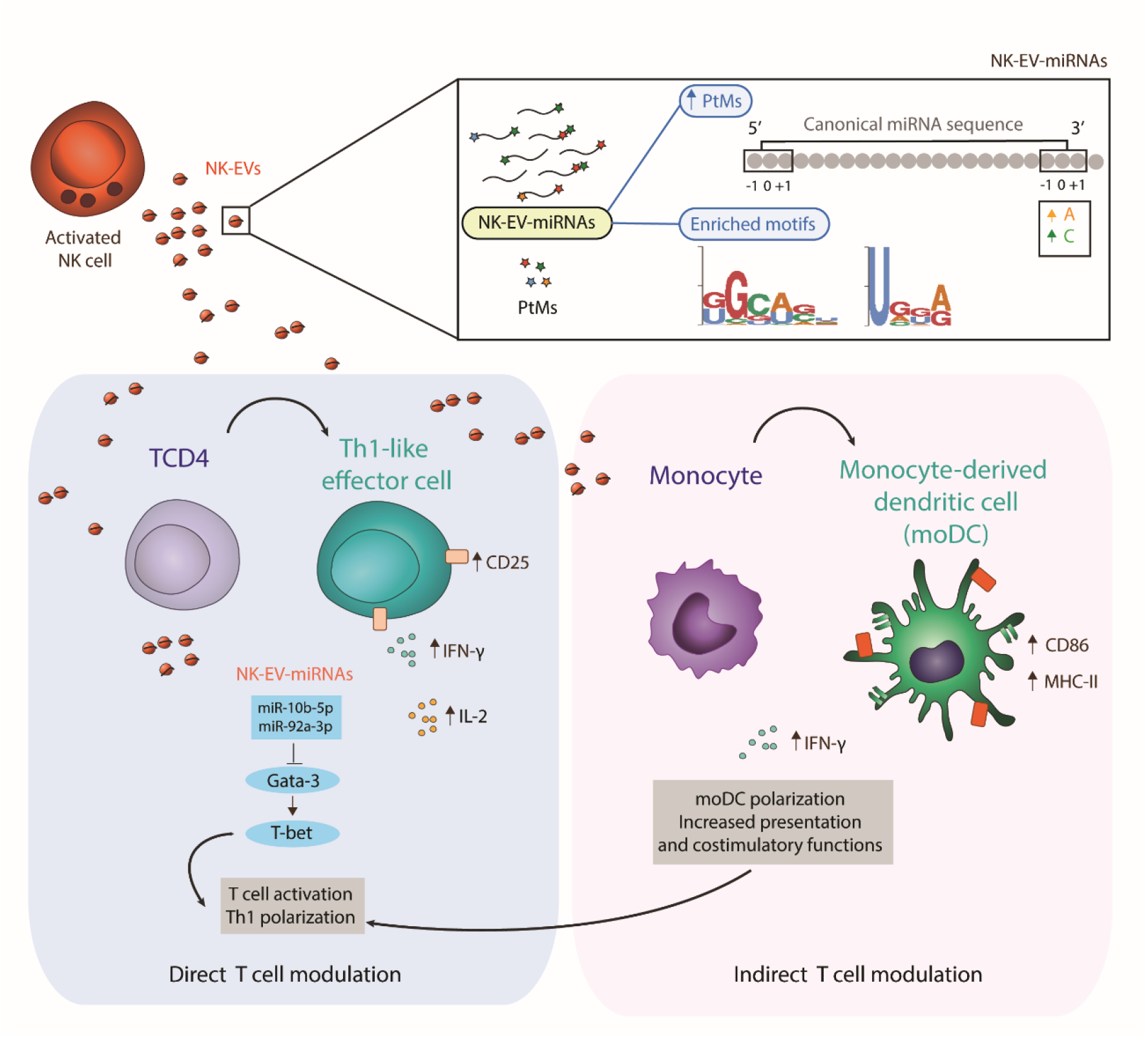
NK-EV miRNAs promote Th1-like responses. NK cells release small EVs that are enriched in post-transcriptionally-modified miRNAs and specific short motifs. NK-EVs have a direct impact on T cells fate and function, promoting Th1-like responses, marked by increased levels of IFN-γ and IL-2 release. NK-EV-miRNAs miR-10b-5p and miR-92a-3p may account, at least partially, for the observed effects, via GATA-3 downmodulation and subsequent T-bet de-repression that drives Th1 skewing. Additionally, NK-EVs promote monocyte polarization to moDCs and enhance their presentation and costimulatory capacities, by upregulating MHC-II and CD86, respectively. This pathway also may contribute to the regulation of Th1-like responses mediated by NK-EVs.

## DISCUSSION

In recent years, the potential of EVs as tools for the treatment, diagnosis and prognosis of several diseases has been intensively studied. Most studies to date suggest that EV miRNA cargo can be exploited to specifically target pathological recipient cells, including immune cells.

NK cells play a pivotal role in the recognition and killing of pathological cells, and recent evidence supports the potential of NK-secreted EVs for therapy (10). Several reports have described the modulation of NK cell effector functions by EVs, including exosomes. Tumour-derived EVs exert immune suppressive functions, dampening lymphocyte cytotoxicity (16, 28, 29). Studies aimed at characterizing the biological effect of dendritic cell (DC)-derived EVs have indicated that they can boost NK and T cell effector functions (30–33). In fact, DC-derived exosomes capabilities are being used in clinical trials for immune maintenance after chemotherapy (34) and cancer immunotherapy (35).

NK-EVs can kill target cells directly (7) due to the presence of cytotoxic proteins amongst the cargo, as shown using proteomics assays (6, 36). Recently, specific NK-EV miRNAs were shown to exert antitumoral effects in neuroblastoma (14) and pancreatic cancer progression (15). In particular miR-186-5p, identified as a down-modulated miRNA in patients with a high risk of developing neuroblastoma was found to inhibit key components of the TGF-β pathway limiting tumour immune evasion. However, the specific NK-EV miRNA repertoire and the impact of NK-EV-miRNAs in the immune response remains far from being fully elucidated.

In this study, we identify a specific miRNA signature of human NK cells (resting and activated) and their secreted EVs by small RNA NGS. Also, we show that a distinct pattern of PtMs is found in resting NK cells, activated NK cells and NK-EVs. Global analysis of miRNA PtMs also identified an enrichment of PtM-miRNAs in the miRNAs more highly expressed in the NK-EV fraction compared to both the relative levels of PtMs in their parental cells, and to the percentage of PtMs found in NK-EVs of activated NK-cell enriched miRNAs. Also, in this context, activated NK cells miRNAs exhibited significantly decreased levels of PtMs compared to resting cells. Thus, it is tempting to speculate that upon activation, post-transcriptionally-modified NK-EV-miRNAs are preferentially targeted to small EVs. These results strongly suggest a PtMs-dependent mechanism of miRNA specific shuttling into NK-EVs.

Analyses of the specific nucleotide additions to the canonical miRNA sequences at different positions, also highlighted a different signature for miRNAs in resting and activated NK cells and their released EVs. The post-transcriptional addition of non-templated nucleotides, particularly to the 3’ends of RNA molecules, is the most frequent and conserved RNA PtM. Interestingly, these modifications have an impact on miRNA stability, and gene regulation, thus on its regulatory capacities (37). Furthermore, 3’uridylation is related to miRNA turnover during T cell activation (38, 39). Non-templated addition of nucleotides has been differentially detected in diverse body fluids. PtMs of miRNAs have been previously reported to determine RNA fate and 3’ adenylation has been found to be overrepresented in cells, while uridylation in exosomes from B cells, suggesting a PtMs-dependent specific cargo of miRNAs into small EVs (40).

In our human NK cell model, we observed that the fraction of both adenylated and cytosylated miRNAs was abundant in resting NK cells and decreased upon activation. Remarkably, a significant enrichment of miRNAs with non-templated 3’ end additions of adenines and cytosines was found in the NK-EV fraction. It is therefore conceivable that cytosylated and adenylated miRNAs are preferentially sorted into small EVs upon NK cell activation.

Moreover, over-represented short sequence motifs were found in NK-EV-miRNAs. In particular, the previously identified GGAG exosomal-sorting motif in T cell-derived small EVs (18) was also enriched in NK-EV-miRNAs, indicating that binding of sumoylated ribonucleoproteins may also be involved in the control of NK-EV-miRNAs sorting. Altogether, our data indicate that a complex regulation, including miRNA PtMs and short motif recognition patterns, may determine the inclusion of specific miRNAs with potential regulatory effects into NK-derived EVs.

Interestingly, NK-EVs are enriched in miRNAs with important functions in the regulation of immune responses, in particular related to Th1 responses. Other miRNAs with immunomodulatory functions are also abundantly expressed in NK cell-derived EVs, e.g. miR-20a-5p and miR-25-3p. These two miRNA are transferred through the immune synapse, with an impact on germinal centre reaction and antibody production (41). Incubation of CD4^+^ T cells with NK-EVs induced an increase of IFN-γ secretion and T-bet expression. This effect correlated with increased levels of the NK-EV associated miR-10b-5p and miR-92a-3p and reduced levels of the T-bet suppressor GATA-3, which is a putative target for these miRNAs (37). In addition, nanoparticle-based delivery of NK-EV-miRNAs miR-92a and miR-155 *in vivo* promoted IFN-γ production. Thus, NK-EV miRNAs miR-10b-5p and miR-92a-3p uptake may promote GATA-3 downregulation and subsequent T-bet de-repression, reprogramming recipient T cells towards the Th1 phenotype. In addition, NK-EVs drive CD4^+^ T cell activation, marked by increased CD25 expression and IL-2 production, but do not significantly affect the Treg compartment. Indeed, although slightly increased levels of Foxp3 were found in T cells after incubation with NK-EVs, no increase in IL-10 secretion was observed and a reduction of this cytokine was found in polarizing conditions, suggesting that NK-EVs may be limiting Treg anti-inflammatory functions in these conditions. A similar trend was observed *in vivo* after nanoparticle-delivery of NK-EV specific miRNAs. In this regard, we detected reduced levels of the pro-inflammatory cytokine IL-6 and TGF-β in T cells upon NK-EV addition. However, the tumour necrosis factor (TNF)-α was found to be increased in T cells shortly after NK-EV addition, in agreement with its presence in the NK92 cell line-derived exosomes (4). It is important to highlight, that EVs do not only contain miRNAs, but also lipids and proteins that also may contribute to the immunomodulatory effects of the NK-EVs. The presence of these non-miRNA bioactive molecules in NK-EVs may account for some of the differences observed between the effects of synthetic miRNAs and NK-EVs.

Our data also show that NK-EVs may impact on T cell responses indirectly by increasing the polarization of monocytes to moDCs, and enhancing their presentation and costimulatory capacities, upregulating MHC-II and CD86 respectively. Hence, NK-EVs promote Th1 responses, both by directly targeting CD4^+^ T cells and by increasing APCs stimulating functions, necessary for the establishment of effective T cells responses. In this regard, many tumours dampen Th effector cell-mediated immunosurveillance, and many immunotherapeutic approaches have focused in restoring Th1 responses (42). Immune therapy has revolutionized the treatment of cancer, however, long-lasting effects are only observed in about 15% of the patients. This might be due to the variety of cancer immune evasion mechanisms, including immune-suppressive mediators in the tumour microenvironment, impairment of CTL responses, promotion of T cell tolerance and/or exhaustion and polarization from Th1 to Th2, among others (43). Thus, we propose that NK-EV mediated immune deviation towards Th1 may be exploited and taken into account when designing EV-based therapies. Interestingly, although primary human NK-EVs isolation may be tedious and time-consuming, a large-scale NK-EV isolation protocol has been set up and may help translation into clinical therapies (44).

In summary, here we provide new insights on the miRNA composition and impact of NK-EVs on immune responses, promoting both directly and indirectly (via APCs) Th1 responses. Unravelling the effects and mechanisms underlying NK-EV mediated immune targeting, in particular Th2 to Th1 immune deviation, may help to improve therapeutic strategies.

## MATERIAL AND METHODS

### In vitro cell culture, antibodies, and reagents

Primary human NK cells were obtained from healthy dono’s buffy coats, using a protocol modified from (45). Briefly, peripheral blood mononuclear cells (PBMCs) were isolated by centrifugation in Ficoll-hypaque and allowed to adhere for 30 min. Thereafter, resting NK cells were obtained using the human NK cell isolation kit (MACS Miltenyi Biotec), following manufacture’s instructions, and lysed in QIAZOL (QIAGEN) buffer for RNA extraction and analysis. For activated NK cultures, PBMCs (seeded at 1,5×10^6^ cells/well) were incubated for 5-6 days in RPMI medium [containing 2 mM L-glutamine, 1 mM sodium pyruvate, 0,1 mM non-essential aminoacids, 100 U/ml Penicillin-Streptomycin (Biowest), 50 μM β-mercaptoethanol (Merck), 10 mM HEPES (Lonza)] supplemented with 10% human serum (HS), 10% foetal bovine serum (FBS), a mixture of irradiated feeder cells (721.221 and RPMI8866, at 0,5×10^6^ per well) and a combination of NK-polarising cytokines; 10 U/ml IL-12 (Peprotech) and 25ng/ml IL-18 (MBL). Then, cells were allowed to proliferate for two additional days by activation with 50 U/ml IL-2 (Peprotech) and 10% HS and 10% FBS. NK cell-enriched cultures were analysed by flow cytometry to confirm the expansion of the NK cell population (CD3^-^CD56^+^). NK cells were purified using the NK isolation kit. Isolated NK cells were cultured for 72 hours in RPMI supplemented with 10% EV-depleted FBS and 50 U/ml IL-2 (by ultracentrifugation at 100,000 g for 16 h) for EV accumulation.

Antibodies and primers are summarized in **Supplementary Table 3**. Primers for specific mature miRNAs detection were purchased from EXIQON-QIAGEN.

### Animals

Wild-type C57BL/6 mice were housed in specific pathogen-free conditions according to European Commission recommendations at the Centro Nacional de Investigaciones Cardiovasculares (CNIC) animal facility in Madrid. Experiments were performed with male mice aged 10–12 weeks. All experimental methods and protocols were approved by the CNIC and the Comunidad Autónoma de Madrid and conformed to European Commission guidelines and regulations.

### NK-EV isolation and characterization

EVs were purified from supernatants after accumulating for 72h from 15×10^6^ NK cells (2×10^6^ cells/ml in RPMI supplemented with 10% EXO-free FBS and 50 U/ml IL-2), by serial centrifugation as previously described (28). EVs were resuspended either in EXO-free RPMI for functional analysis, PBS for EV characterization by NanoSight (41), or in Laemmli loading buffer for western blot.

For small RNA sequencing and qPCR analysis, EV pellets obtained after 100,000xg ultracentrifugation of NK cell supernantants, were lysed in QIAZOL (QIAGEN) and RNA was isolated using the miRNeasy Mini Kit, as described above. Culture medium pellet after 100,000g ultracentrifugation alone was analysed by size-exclusion chromatography to rule out possible serum contaminations in NK-EV samples, as previously described (41).

All the relevant data of our experiments were submitted to the EV-TRACK knowledgebase (EV-TRACK ID: EV210234) (46).

### RNA extraction, real-time PCR, small RNA next-generation sequencing (NGS), and data analysis

RNA from purified human NK cells, both resting and activated *in vitro* (as described above) and their secreted small EVs were extracted using the miRNeasy Mini Kit (QIAGEN).

For small RNA sequencing, RNA integrity was checked using an Agilent 2100 Bioanalyzer (Agilent) for total RNA (RNA nano-chips) and for small RNA (small RNA chips), and concentrations were measured in a Nanodrop-1000 and using the Quantifluor RNA system (Promega). Samples from 5 healthy donors were analysed, and small RNA libraries were generated using the NEBNext Small RNA Library Prep Set from Illumina. Single read NGS was performed using an Illumina HiSeq 2500 System. Small RNASeq data were analysed by the Bioinformatics Unit at CNIC. Sequencing reads were processed as previously described (41). Changes in small RNA expression were considered significant when Benjamini and Hochberg adjusted P-value < 0.05.

For qPCR analysis, isolated RNA was retro-transcribed using either the miRCURY LNA Universal RT miRNA PCR System (EXIQON) for miRNA or the Promega RT kit for mRNA, as described in (41). Real-time (RT)–PCR was performed in a CFX384 Real-time System (Bio-Rad) using SYBR Green PCR Master Mix (Applied Biosystems). Reactions were analysed with Biogazelle QbasePlus software. Mature miRNA levels were normalized to the small nucleolar RNAs RNU5G and RNU1A1, whereas mRNA levels were normalized to glyceraldehyde-3-phosphate dehydrogenase (GAPDH) and β-actin and expressed as a relative variation from control levels. All primers for miRNA analysis were purchased from EXIQON, and primers used for mRNA detection (Metabion) are listed in supplementary methods.

### In silico target analysis

Putative mRNA targets for human miRNAs overexpressed in small EVs compared to secreting NK cells, were identified using the prediction algorithm miRTarBase and are summarized in **Supplementary Table 2**.

Functional unbiased analysis of EV-enriched miRNAs (compared to secreting NK cells) was performed using the Ingenuity Pathway Analysis (IPA).

### NK-EVs functional assays

#### NK-EVs and CD4+ T cell function assays

CD4^+^ T cells were isolated from non-adherent PBMCs, using the human CD4^+^ T cell isolation kit (MACS Miltenyi Biotec). Isolated T helper cell purity was checked by flow cytometry CD4 staining and they were then cultured in CD3 (5µg/ml) and CD28 (2µg/ml) coated plates either in the presence or absence of NK-EVs and both in non-polarizing (RPMI supplemented with 10% EV-free FBS) and Th1 polarizing conditions, adding a mixture of cytokines (20U/ml IL-2 and 20ng/ml IL-12). NK-EVs obtained from 15×10^6^ NK cells supernatants, as described above, were resuspended in 100µl of EXO-free RPMI and 25µl were added to 2×10^6^ CD4^+^ T cells per condition. On days 3 and 6 post-coculture, cells in the different conditions and their supernatants were analysed as described below. NK-EVs isolated from three different healthy donors were cocultured with Th cells from at least two different donors.

### NK-EVs and monocyte/moDC function assays

Human monocyte–derived DCs (moDCs) were obtained as described (26). Briefly, CD14^+^ monocytes were isolated from total PBMCs, using a positive selection kit (StemCell) and cultured in RPMI medium supplemented with 10% EV-free FBS, either in the presence or absence of DC-polarizing cytokines (50ng/ml GM-CSF and 1000U/ml IL-4). The effects of NK-EVs addition (25µl of isolated NK-EVS were used to treat 3×10^6^ CD14^+^ cells) were analysed 6 days after culture. NK-EVs isolated from three different healthy donors were cocultured with CD14^+^ cells from at least two different donors.

### Flow cytometry

Cultured cells were stained with the antibodies listed in **Supplementary Table 3** and analysed by flow cytometry in a BD FACSCanto cytometer. For all analysis, only live single cells were included and analysed with the FlowJov10 software. Dead cells were excluded by DAPI or Live/Dead Fixable Yellow (Thermofisher) staining. Murine cells were incubated with mouse CD16/CD32/Fc shield and human cells with 100µg/ml γ -globulin before antibody staining.

### Cytokine Arrays

CD4+ T cells were isolated as described above from human buffy coats. A total of 5×10^6^ cells were incubated for 16 hours with NK-EVs isolated from 5×10^6^ 72 h NK cell cultures in RPMI supplemented with 10% EV-free FBS. Culture supernatants were analysed using the Proteome Profiler Human Cytokine Array Kit (R&D Systems), following manufacture’s instructions.

### Cytokine analysis of cell culture supernatants by ELISA

Cytokine production was detected in cell culture supernatants using ELISA kits purchased from DIACLONE (IL-2) and eBioscience (IFN-γ) respectively, following manufacture’s instructions. Absorbance was measured at 450 nm, with a reference wavelength of 570 nm.

### Nanoparticle-miRNAs synthesis

Gold nanoparticles (AuNPs) were synthesized following the Turkevich method (47). Rounded 12 nm AuNPs at 25.2 nM were obtained and modified with oligonucleotides (IDT) containing the proper thiol-modifications **(Supplementary Table 4)**. MiRNA duplex formation was performed using the same volume and concentration of both strands diluted in annealing buffer (10 mM Tris, pH 7.5 - 8.0, 50 mM NaCl, 1 mM EDTA), according to Sigma Aldrich’s protocol (https://www.sigmaaldrich.com/ES/es/technical-documents/protocol/genomics/pcr/annealing-oligos). The mixture was incubated for 10 min at 95 °C and then cooled down to RT slowly. Before conjugation, the miRNA duplexes were deprotected by incubation with 100x excess TCEP for 2h at RT. For the functionalization reaction, 4 pmol/ µl AuNPs of deprotected miRNA duplex were added to 1 ml AuNP for each condition. AuNP miR-mix contains 4 pmol/ µl of each miRNA duplex; AuNP control contains 4 pmol/ µl poli T-thiol (TTTTTTTTTTTT/3ThioMC3-D/). After oligonucleotide addition, NaCl was added in small volumes until 0.3M final concentration. Finally, the AuNPs were incubated 16 h in agitation, light protected at RT. After this step, 3 washeswere performed by 40 min 16,100 g centrifugation at 4°C. In the last washing step, AuNPs were resuspended in sterile 1x PBS and filtered with 0.2 µm syringe-filter in a laminar hood.

### Nanoparticle-miRNAs analysis

Nanoparticles in PBS were injected into the footpad (30µl) of wild-type C57BL/6 mice. On day 5, animals were sacrificed and splenocytes isolated and analysed by flow cytometry and qRT-PCR before and after stimulation with anti-CD3 and anti-CD28 antibodies. All cells were incubated with PMA (50ng/ml) and ionomycin (500ng/ml) 4h before analysis, and Brefeldin A (5µg/ml) was added to block secretion in flow cytometry experiments. Supernatants from cultured cells after 16h culture were also analysed by ELISA.

### Small RNASeq analyses

Small RNASeq data were analyzed by the Bioinformatics Unit at CNIC. Sequencing reads were pre-processed by means of a pipeline that used FastQC (http://www.bioinformatics.babraham.ac.uk/projects/fastqc/), to asses read quality, and Cutadapt to trim sequencing reads, eliminating Illumina adaptor remains, and to discard those that were shorter than 20 nucleotides after trimming. Around 60% of the reads from any of the samples were retained. Resulting reads were aligned against a collection of 2657 human, mature miRNA sequences extracted from miRBase (release 21), to obtain expression estimates with RSEM (38). Percentages of reads participating in at least one reported alignment were around 22%. Expected expression counts were then processed with an analysis pipeline that used Bioconductor package Limma (48) for normalization (using TMM method) and differential expression testing, considering only 708 miRNA species for which expression was at least 1 count per million (CPM) in 3 samples. Changes in gene expression were considered significant if the Benjamini-Hochberg adjusted p-value < 0.05.

### miRNA PtMs analyses

Epi-transcriptomic modifications were detected with Chimira (49), an online tool that, after alignment of miRNA-Seq reads against miRBAse21 records, identifies mismatched positions to classify them and to quantify multiple types of 3’-modifications (uridylation, for example), as well as internal and 5’-modifications. Count tables produced by Chimira were further processed with ad-hoc produced R-scripts to calculate summary statistics across groups of replicate samples.

### Motif enrichment analysis

Over represented motifs in miRNA collections were detected with MEME (50), using the differential enrichment method and complementary miRBAse21 miRNA collections as background references. Both types of input were previously processed with USEARCH to make them non redundant at 80% identity (51). Motif searches were performed only in the forward strand, using a motif size of 4-8 nt, 0-order Markov model for nucleotide distributions and expecting zero or one motif occurrence per sequence (zoops).

### Statistical analyses

All data were analysed with GraphPad Prism 5.0 and 8.0 (GraphPad Software, San Diego, CA, USA). All data were included in the analysis, unless identified as outliers, using GraphPad algorithms. Appropriate statistical tests were used, for each experiment, as indicated. Significance was set at *p < 0.05, **p < 0.01, ***p < 0.001.

### Data availability

Sequencing data have been deposited in the Gene Expression Omnibus and are available to readers under record GSE185171 (Password: szgtqawoxpwvhit). EV isolation procedures are available at EV-TRACK knowledgebase (EV-TRACK ID: EV210234.

## ACKNOWLEDGEMENTS

NGS experiments were performed in the CNIC Genomics Unit (Centro Nacional de Investigaciones Cardiovasculares, Madrid, Spain) and analysed by the CNIC Bioinformatics Unit. This manuscript was funded by grants PD1-2020-120412RB-100 (FS-M) and PID2020-119352RB-I00 (AS) from the Spanish Ministry of Economy and Competitiveness; CAM (S2017/BMD-3671-INFLAMUNE-CM) from the Comunidad de Madrid (FS-M). CIBERCV (CB16/11/00272) and BIOIMID PIE13/041 from the Instituto de Salud Carlos. The current research has received funding from “la Caixa” Foundation under the project code HR17-00016. Grants from Ramón Areces Foundation “Ciencias de la Vida y de la Salud” (XIX Concurso-2018) and from Ayuda Fundación BBVA y Equipo de Investigación Científica (BIOMEDICINA-2018) (to FSM). The CNIC is supported by the Ministerio de Ciencia, Innovacion y Universidades and the Pro-CNIC Foundation, and is a Severo Ochoa Center of Excellence (SEV-2015-0505). IMDEA Nanociencia acknowledges support from the ‘Severo Ochoa’ Programme for Centres of Excellence in R&D (MINECO, CEX2020-001039-S). SGD is supported by a grant from the Spanish Ministry of Universities. Authors thank Dr. Miguel Vicente-Manzanares for critical review and editing.

## AUTHOR CONTRIBUTIONS

LFM planned and performed experiments, analysed and interpreted data, and wrote the manuscript with input from all authors; SGD and SLC helped with research design and NK cell cultures and exosome isolation experimentation; SGD, ARG and IFD contributed with functional assays experimentation and mice models; MC, AS and PMR designed and generated miR-nanoparticles; MVG and HTR helped with research design, provided reagents, and collaborated in data interpretation and manuscript writing; FSM planned and coordinated the research, discussed results, and supervised and contributed to manuscript writing.

## SUPPLEMENTARY MATERIALS

**Fig. S1.**
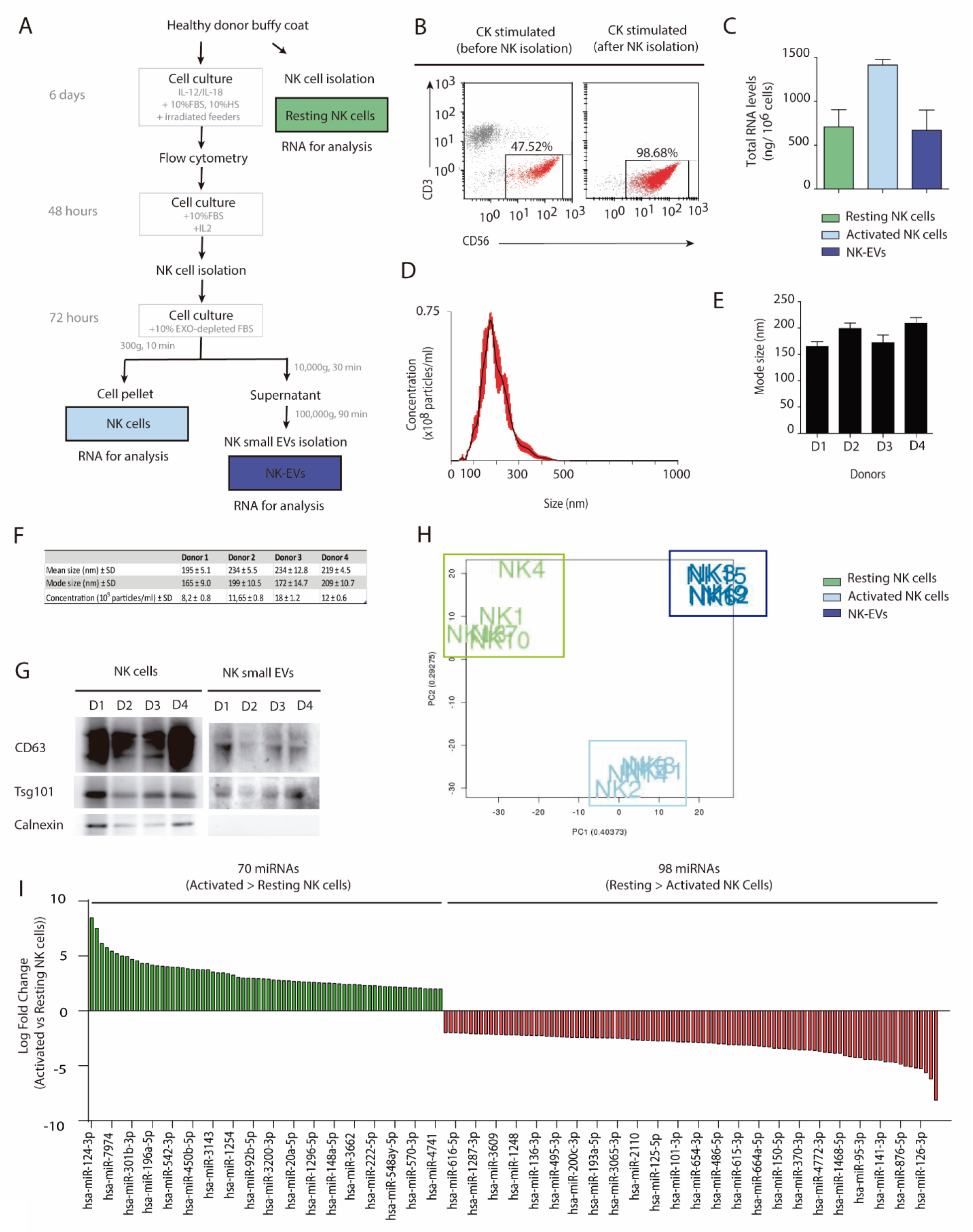
NK-EV isolation and characterization. (**A**) Scheme of NK cells (resting and activated) and NK-EV RNA samples preparation for small RNA sequencing analysis. (**B**) Representative flow cytometry dot plot of PBMCs from healthy dono’s buffy coats in the presence of NK-enrichment cytokine (CK) cocktail with IL-12 and IL-18 for 6 days. Dot plots show the enrichment of CD3-CD56+ NK cells after 6-8 days culture with cytokines and irradiated feeder cells (left panel) and NK cell isolation (right panel), being greater than 95% in all experiments. (**C**) Quantification of total RNA (Nanodrop) from isolated samples for small RNA sequencing analyses (resting and activated cells and NK-EVs). (**D-F**) Characterization of isolated NK-derived small EVs by NanoSight analysis. A representative plot from all analysed samples is shown (**D**). Nanosight determination of size quantification (**E**) and statistics, including mean and mode size and EV concentration (**F**) of EVs isolated from 4 healthy donors, as described in the manuscript’s methods section. (**G**) NK cell total lysates and NK-EV fractions were analysed by Western blot, probing with the exosomal markers Tsg101 and CD63 and Calnexin, as indicated. (**H**) Principal Component Analysis plot from small RNA sequencing data; resting NK cell (green), activated NK cells (light blue) and exosome samples (dark blue) are represented. (**I**) Waterfall plot showing logarithmic fold-increase expression between resting and activated NK cells. Only fold-changes with adjusted P-value < 0.05 are represented. MiRNAs significantly increased in activated NK cells are shown in green, while miRNAs significantly more represented in resting cells are shown in red.

**Fig. S2.**
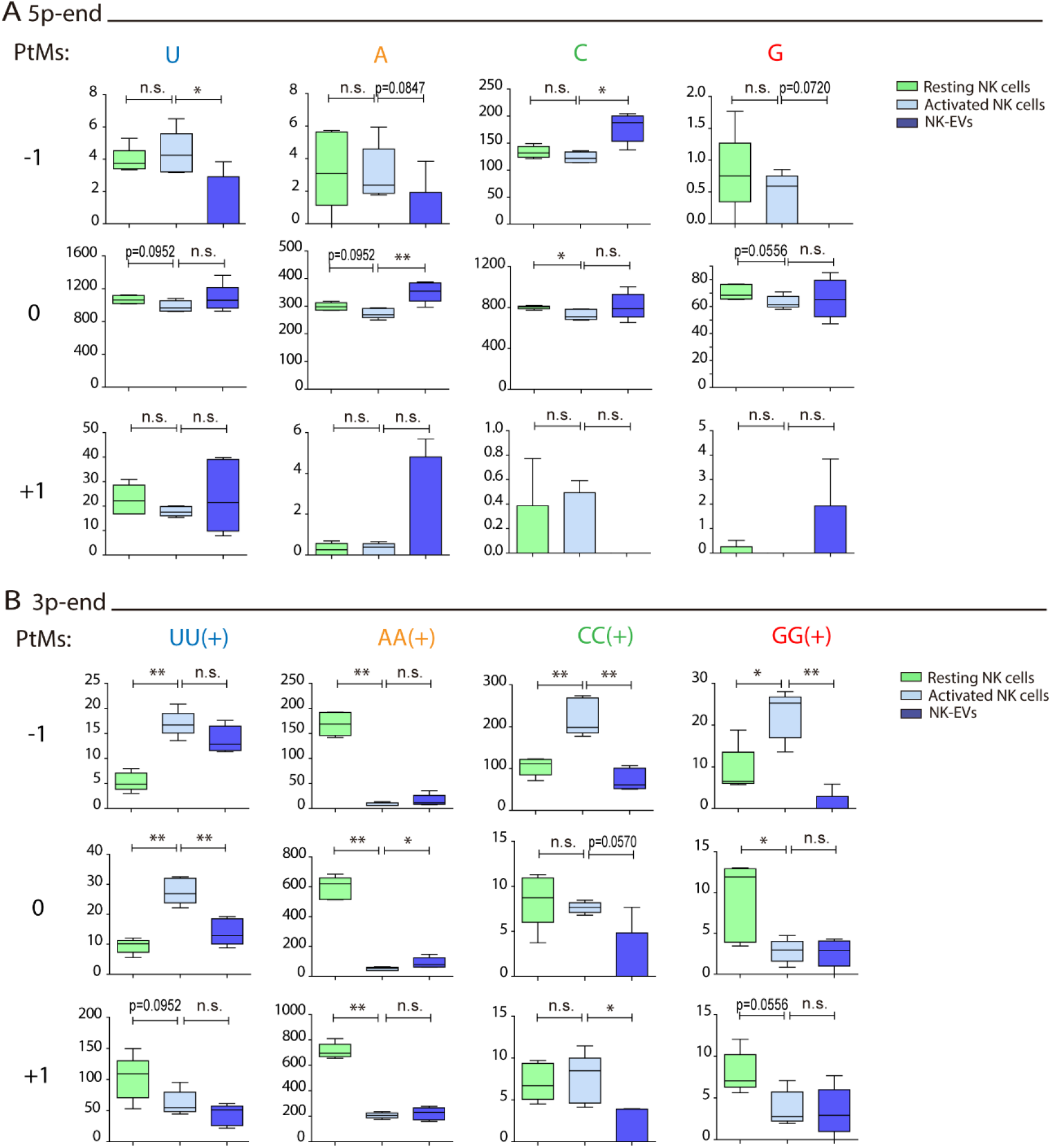
Resting NK cells, activated NK cells and NK-EV miRNAs have a different PtM signature. (**A**) Box and whiskers plots show the mono-additions of U, A, C and G at the indicated positions (ranging from −1 to +1) in miRNAs from resting NK cells, activated NK cells and NK-EVs at the 5p-end. Significance was assessed by non-parametric t-test, comparing resting with activated NK cells, and activated NK cells with their released NK-EVs, respectively; *P<0.05, **P<0.01. (**B**) Box and whiskers plots show the additions of two or more U, A, C and G at the indicated positions (ranging from −1 to +1) in miRNAs from resting NK cells, activated NK cells and NK-EVs at the 3p-end. Significance was assessed by non-parametric t-test, comparing resting and activated NK cells and activated NK cells and their released NK-EVs, respectively; *P<0.05, **P<0.01.

**Fig. S3.**
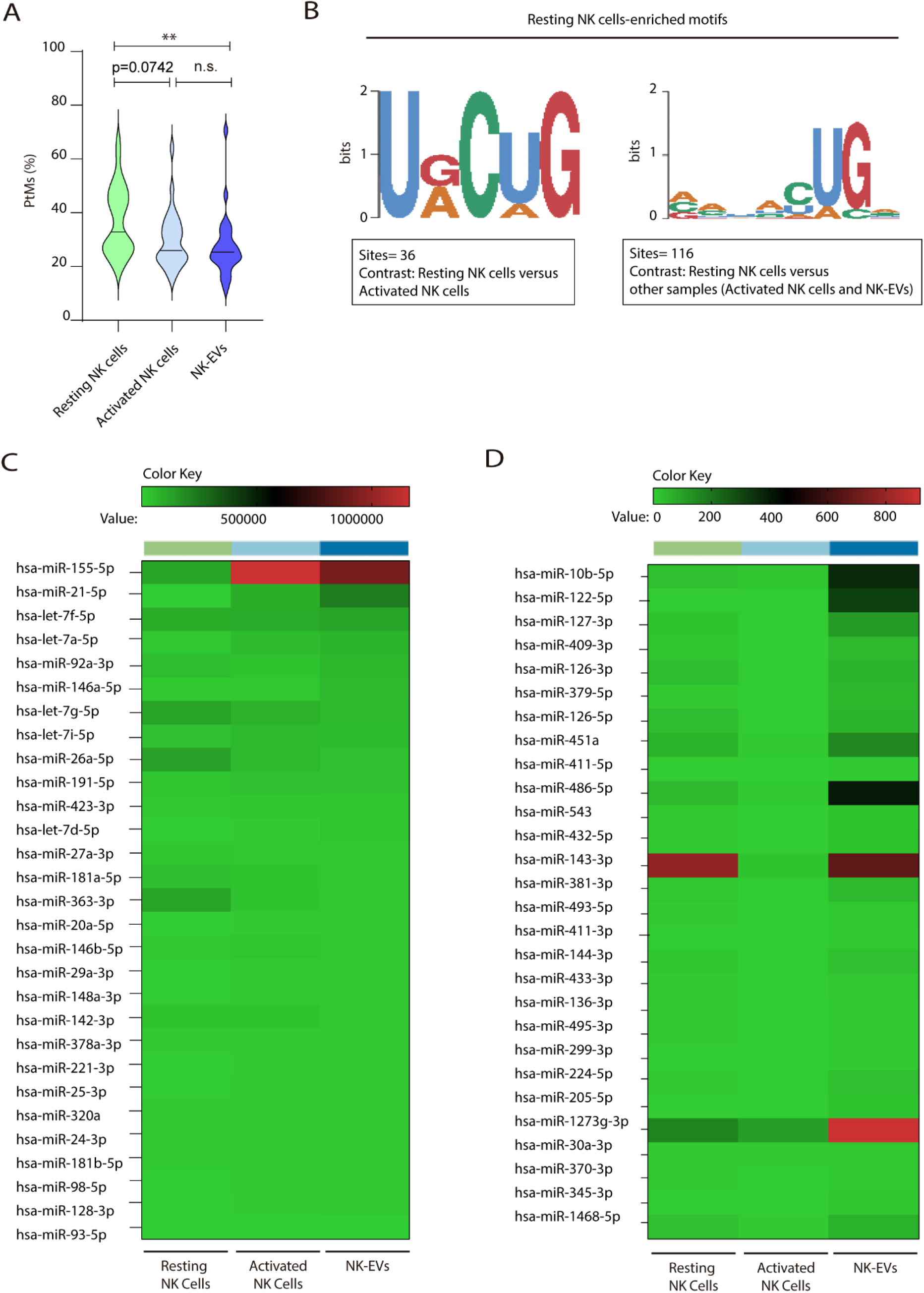
PtM characterization of the most expressed miRNAs in NK-EVs and resting NK cells over-represented motifs. (**A**) Violin plots represent the average percentage of reads with PtMs from each miRNA. Analysis was performed comparing the most expressed miRNAs in NK-EVs. The median value is shown as a solid line. (**B**) Over-represented motifs in resting NK cells are shown, using the ZOOPS model. For each data set, all miRNAs annotated in miRBase were used as background. Short motifs with adjusted E-value<0.2 are shown. (**C,D**) Heat maps showing small RNA sequencing analysis of the top 30 miRNAs with higher levels of expression in NK-EVs (**C**) and those more differentially expressed in NK-EVs compared to activated NK cells (**D**), respectively. For each analysis, the relative expression of individual miRNAs in resting and activated human NK cells and their secreted exosomes is shown. Data are from 5 healthy dono’s isolated NK cells.

**Fig. S4.**
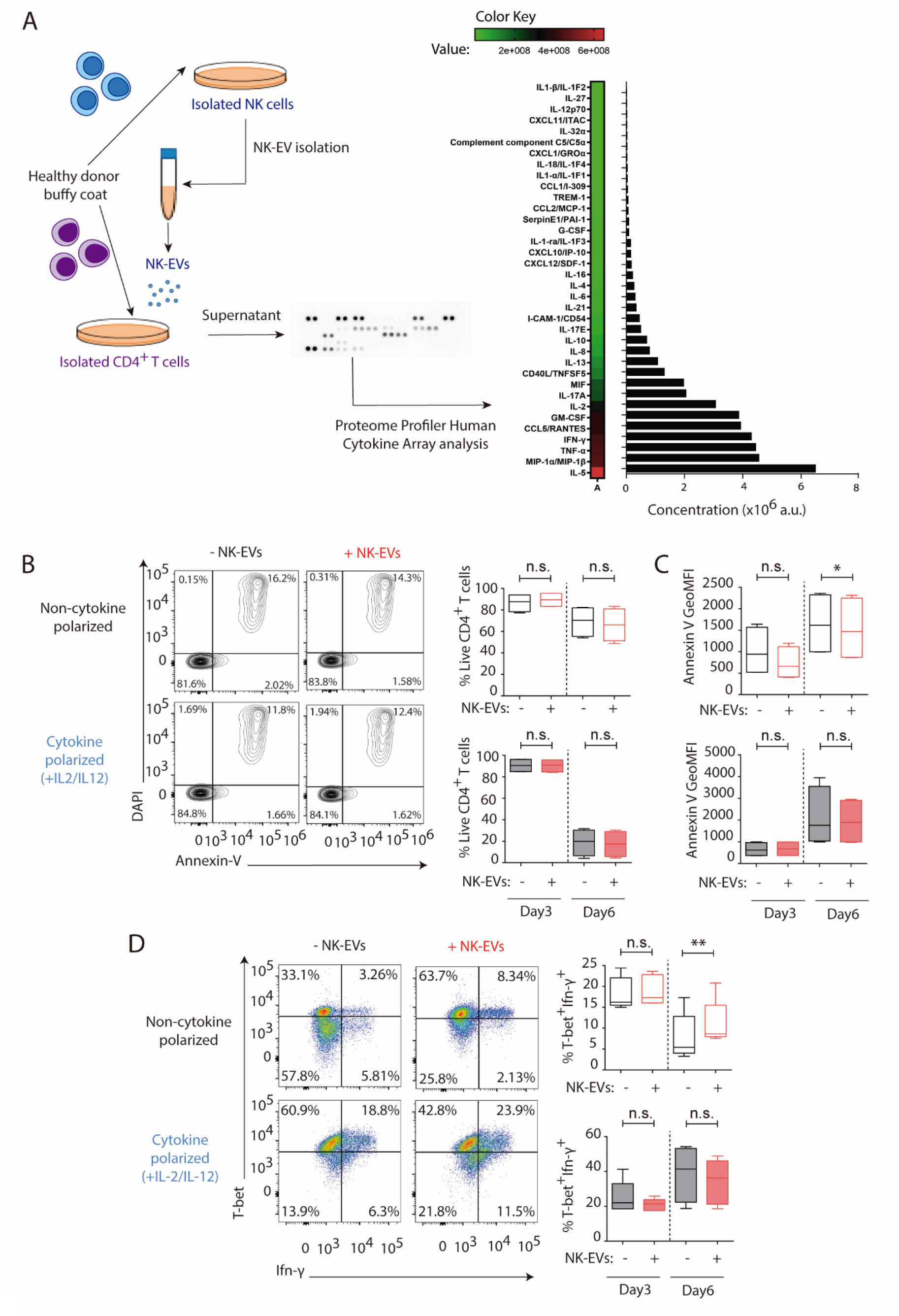
NK-EV miRNAs and T cell function. **(A)** Cytokine profiler of CD4+T lymphocytes incubated for 16 h with NK-EVs. Data show the relative expression of the indicated cytokines and chemokines, expressed in arbitrary units (a.u.). (**B**) Flow cytometry analysis showing the viability of isolated CD4+ T cells incubated under non-cytokine polarizing (upper panel) and Th1 cytokine-polarizing (with a mixture of IL-2 and IL-12, lower panel) conditions. Contour plots show a representative staining with the viability marker DAPI and expression of the apoptotic marker Annexin-V in CD4+ T cells. Plots show the quantification of live DAPI-Annexin-V- CD4+ T cells ± SEM, in the different culture conditions and after addition of NK-EVs. Plots show the quantification of n ≥ 4 independent experiments. Significance was assessed with Paired Student’s t-test. (**C**) Quantification of Annexin-V Geometric Mean Fluorescence Intensity. Data shown are from n ≥ 4 independent experiments. Significance was assessed with Paired Student’s t-test; *P<0.05. (**D**) Flow cytometry analysis of isolated CD4+ T cells incubated under non-cytokine polarizing (upper panel) and Th1 cytokine-polarizing (with a mixture of IL-2 and IL-12, lower panel) conditions. Dot plots show the expression of T-bet and IFN-γ in gated live cells ± SEM, after addition of NK-EVs. Plots show the quantification of n ≥ 4 independent experiments. Significance was assessed with Paired Student’s t-test; *P<0.05, **P<0.01.

**Fig. S5.**
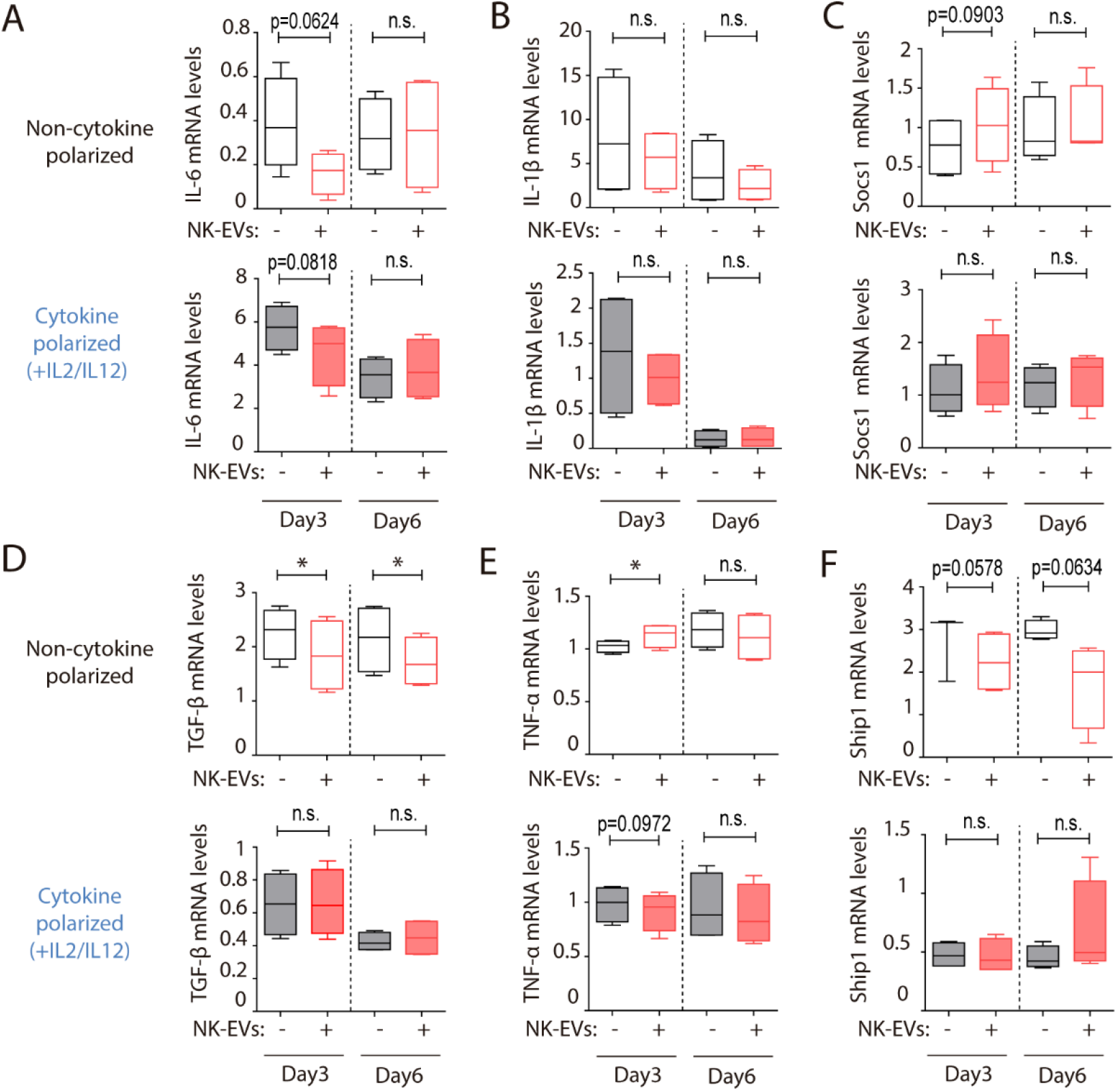
mRNA target modulation in CD4+ T cells mediated by NK-EVs. (**A-F**) Quantitative real-time PCR at days 3 and 6, as indicated in either non-polarizing (upper panels) or cytokine-polarizing (lower panels) conditions, showing mRNA levels of IL-6 (**A**), IL-1β (**B**), Socs1 (**C**), TGF-β (**D**), TNF-α (**E**) and Ship1 (**F**) respectively, normalized to GAPDH and β-actin. Significance was assessed with Paired Student’s t test; *P<0.05.

**Fig. S6.**
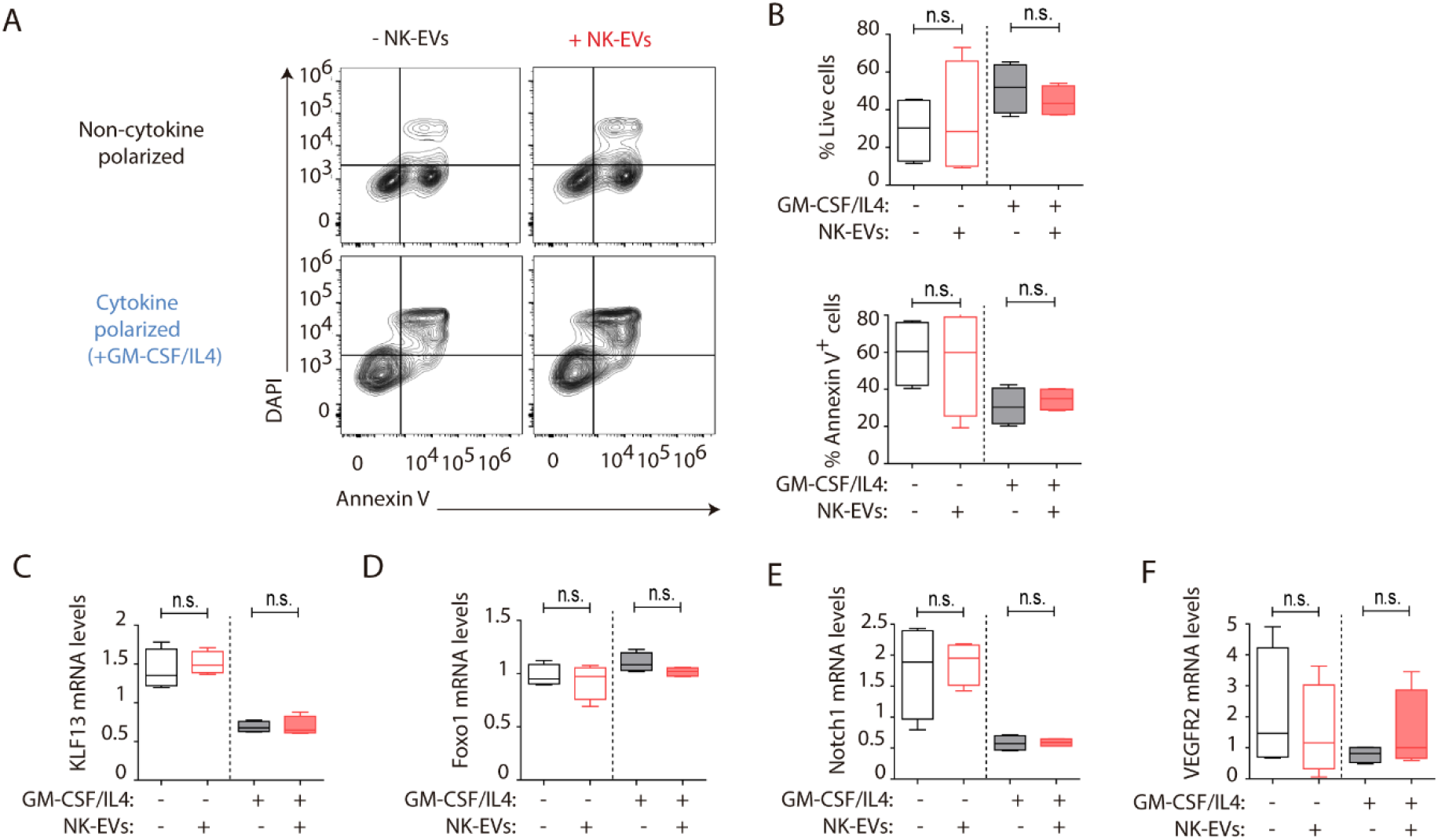
NK-EVs impact on monocyte and moDC function. (**A**) Flow cytometry viability analysis of isolated CD14+ monocytes (non-polarized, upper panel) and moDCs (cytokine-polarized with a mixture of GM-CSF and IL-4, lower panel) in the indicated culture conditions. Contour plots show a representative staining with the viability marker DAPI and the apoptotic marker Annexin-V in cultured cells. (**B**) Plots show the quantification of live DAPI-Annexin-V-CD14+ monocytic cells ± SEM, in the different culture conditions and after addition of NK-EVs. Plots show the quantification of n ≥ 4 independent experiments. Significance was assessed with Paired Student’s t-test. (**C-F**) Quantitative real-time PCR at day 6, of isolated CD14+ monocytes, cultured as indicated in either non-polarizing or cytokine-polarizing conditions. Plots show the relative mRNA levels of the putative NK-EV miRNAs mRNA targets KLF13 (**C**), Foxo-1 (**D**), Notch1 (**E**) and VEGFR2 (**F**) respectively, normalized to GAPDH and β-actin. Significance was assessed with Paired Student’s t test.

**Fig. S7.**
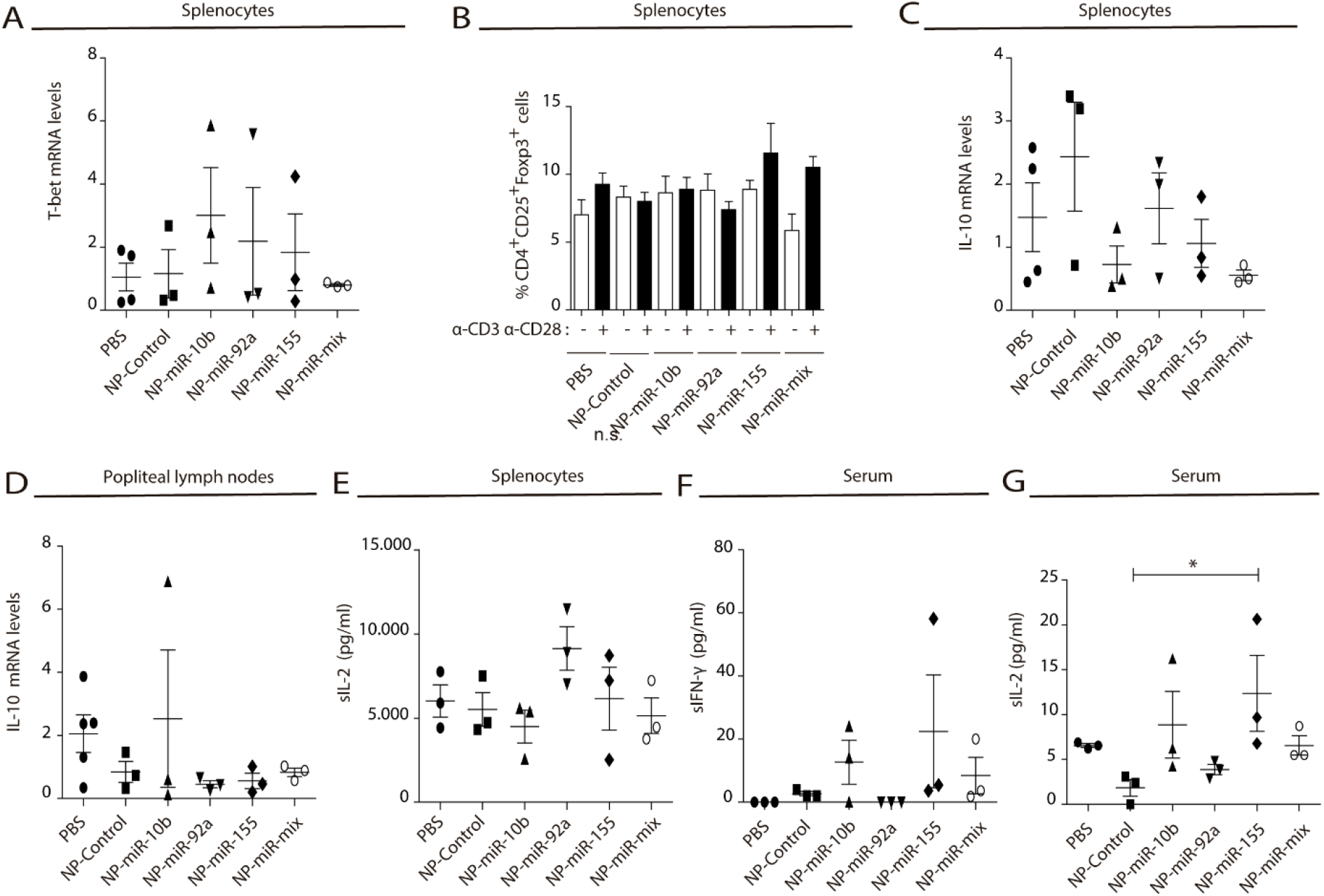
NK-EV miRNAs partially mimic NK-EV mediated effects in vivo in spleen and draining popliteal lymph nodes. (**A**) Quantitative real-time PCR showing T-bet mRNA levels in splenocytes 6 days after nanoparticle-based miRNA delivery. Relative expression is shown, normalized to GAPDH and β-actin. Significance was assessed by One-way ANOVA Bonferroni test. (**B**) Flow cytometry analysis of live splenocytes isolated 6 days after footpad injection with the indicated control or miRNA-loaded NPs. Isolated cells were either left unstimulated (white) or incubated for 16 h with anti-CD3 and anti-CD28 antibodies (black) before analysis. Bars show the percentage of CD4+CD25+Foxp3+ live T lymphocytes. Experiments from n ≥ 3 mice are shown. Significance was assessed by Two-way ANOVA, followed by Tukeýs test. (**C,D**) Quantitative real-time PCR showing IL-10 mRNA levels in splenocytes (**C**) and in popliteal draining lymph nodes (**D**) 6 days after nanoparticle-based miRNA delivery. Relative expression is shown, normalized to GAPDH and β-actin. Significance was assessed by One-way ANOVA Bonferroni test. (**E**) ELISA quantification of soluble IL-2 in supernatants from splenocytes harvested and isolated after 6 days miRNA delivery and cultured with anti-CD3 and anti-CD28 antibodies for 16 h. The graph shows the mean concentration from n ≥ 3 mice per condition. Significance was assessed by One-way ANOVA Bonferroni test. (f, g) ELISA quantification of soluble IFN-γ (**F**) and IL-2 (**G**) in serum from mice 6 days after NP footpad injection. The graph shows the mean concentration from n ≥ 3 mice per condition. Significance was assessed by One-way ANOVA Bonferroni test; *P<0.05.

**Table S1.** Small RNA sequencing of resting NK and activated NK cells and their derived NK-EVs. Table summarizes the analysis of samples generated from 5 donors, showing differentially expressed miRNAs. Provided as an additional Excel file.

**Table S2.** NK-EV miRNAs mRNA target candidates. Table summarizes putative mRNA targets for selected NK-EV miRNAs identified by in silico analyses, using the miRTarBase database. T cell function-related miRNAs are highlighted in orange and DC related miRNAs in blue.

| miRNA | Fold-change | Target | Function | Reference |
| --- | --- | --- | --- | --- |
| miR-155 | 2.51 | Ship1<br>Socs1 | Activation T cell and clonal selection<br>DC maturation<br>MHC-II, costimulatory molecules, IL12, TNF- and DC-sign upregulation | Baumjohann et al, Nat, 2013<br>Turner ML et al., JI, 2016 |
| miR-10b | 806.11 | GATA3<br>PTEN | De-repress T-bet Th1 response,<br>Inhibit Th2 | mirTarbase |
| miR-1-3p | 6.58 | GATA3 | De-repress T-bet Th1 response | mirTarbase |
| miR-92a-3p | 1.46 | GATA3 | De-repress T-bet Th1 response | mirTarbase |
|  |  | Pten<br>Bim | T cell survival and proliferation | Baumjohann et al, Nat, 2013 |
| miR-335-5p | 3.32 | STAT6 | De-repress T-bet Th1 response,<br>Inhibit Th2 | mirTarbase |
| let-7b | 2.39 | Cox2? | Prevention of systemic disease by suppression of pathogenic Th1 | Okoye IS, Immunity, 2014 |
| let-7d-5p | 2.60 | Cox2? | Prevention of systemic disease by suppression of pathogenic Th1 | Okoye IS, Immunity, 2014 |
| miR-146a-5p | 3.32 |  | Maturation/activation DCs<br>Survival/maturation pDCs | Turner ML et al., JI, 2016 |
| Let-7c | 8.54 | Blimp | Maintaining tolerogenic DCs | Sun Jung Kim. JCI, 2013 |
| miR-222-3p | 1.71 |  | Maturation cDCs | Turner ML et al., JI, 2016 |
| miR-370-3p | 14.75 | FOXO1 | Predict target FOXO1 inhibits via NF-kB apoptosis of DCs | Riol-Blanco L, Nat Immunol, 2009<br>miRTarbase |
| miR-451a | 79.75 | Notch1 | Tunes DC cytokine production | Lesley A Smyth, Immunology 2015 |
| miR-126-3p | 89.43 | VEGFR2 | Survival and function of pDCs<br>Regulates expression of molecules from IIR | Agudo j, Nat Immunol, 2014 |
| mir-99a-5p | 4.71 | KLF13 | Promotes tolerogenic DCs phenotype and function | Lesley A Smyth, Immunology 2015 |

**Table S3.**
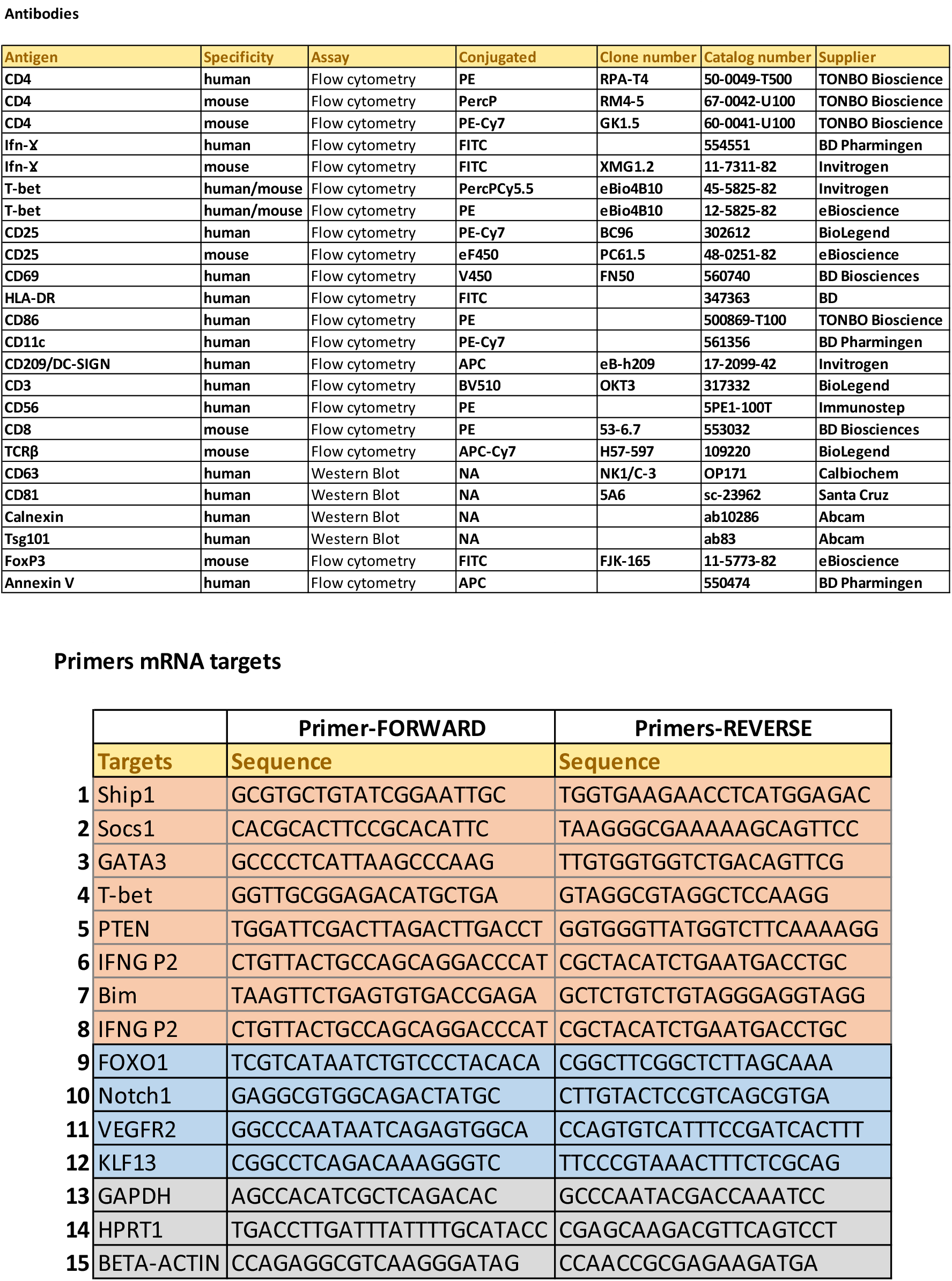
Antibodies and primers. Specific primers for the detection of mRNA related with T-cell functions are shown in orange and DC-function related miRNAs in blue. Housekeeping mRNAs are shown in gray.

**Table S4.** Oligonucleotides for miRNA duplex AuNPs design.

|  | miR-10b-5p | miR-92a-3p | miR-155-5p |
| --- | --- | --- | --- |
| miRNA<br>guide<br>strand | rUrArCrCrCrUrGrUr<br>ArGrArArCrCrGrArA<br>rUrUrUrGrUrG | rUrArUrUrGrCrArCrUrUr<br>GrUrCrCrCrGrGrCrCrUr<br>GrU | rUrUrArArUrGrCrUrArA<br>rUrCrGrUrGrArUrArGr<br>GrGrGrUrU |
| miRNA<br>pass<br>strand | rCrArCrArArArUrUrC<br>rGrGrUrUrCrUrArCr<br>ArGrGrGrUrArUrUrU<br>rUrU/3ThioMC3-D/ | rArCrArGrGrCrCrGrGrGr<br>ArCrArArGrUrGrCrArArU<br>rArUrUrUrUrU/3ThioMC3<br>-D/ | rArArCrCrCrCrUrArUrC<br>rArCrGrArUrUrArGrCrA<br>rUrUrArArUrUrUrUrU/3<br>ThioMC3-D/ |

